# A repurposed, non-canonical cytochrome *c*, chaperones calcium binding by PilY1 for type IVa pili formation

**DOI:** 10.1101/2021.07.28.454143

**Authors:** Marco Herfurth, Anke Treuner-Lange, Timo Glatter, Nadine Wittmaack, Egbert Hoiczyk, Antonio J. Pierik, Lotte Søgaard-Andersen

## Abstract

Type IVa pili (T4aP) are versatile bacterial cell surface structures that undergo extension/adhesion/retraction cycles powered by the cell envelope-spanning T4aP machine. In this machine, a complex composed of four minor pilins and PilY1 primes T4aP extension and is also present at the pilus tip mediating adhesion. Similar to many other bacteria, *Myxococcus xanthus* contains multiple minor pilins/PilY1 sets that are incompletely understood. Here, we report that minor pilins and PilY1 (PilY1.1) of cluster_1 form priming and tip complexes contingent on a non-canonical cytochrome *c* (TfcP) with an unusual His/Cys heme ligation and calcium. We provide evidence that TfcP is unlikely to participate in electron transport and has been repurposed to promote calcium binding by PilY1.1 at low calcium concentrations, thereby stabilising PilY1.1 and enabling T4aP function in a broader range of calcium concentrations. These results identify a novel function of cytochromes *c* and illustrate how incorporating an accessory factor expands the environmental range under which the T4aP system functions.

## Introduction

In bacteria, motility is important for virulence, promotes colonisation of habitats of diverse composition, and stimulates biofilm formation^1^. Type IVa pili (T4aP) are filamentous cell surface structures that enable cell translocation across surfaces and also have critical functions in surface adhesion, surface sensing, host cell interaction, biofilm formation, predation, virulence, and DNA uptake^2–4^. The versatility of T4aP is based on their ability to undergo cycles of extension, surface adhesion, and retraction^5, 6^. Retractions generate a force up to 150 pN per pilus and pull cells across surfaces^7^.

In Gram-negative bacteria, the extension/retraction cycles of T4aP are driven by the T4aP machine (T4aPM), which consists of 15 conserved proteins that form a complex that spans from the outer membrane (OM) across the periplasm and inner membrane (IM) to the cytoplasm^8–10^ (Fig. 1a). Pilus extension and retraction are powered by the PilB and PilT ATPases, respectively, that bind in a mutually exclusive manner to the cytoplasmic base of the T4aPM^8, 11–13^. All 15 proteins are essential for T4aP extension except for PilT, which is only important for retraction^4^. The so-called priming complex is an integral part of the T4aPM, is composed of the major pilin, four minor pilins and the PilY1 protein, and is incorporated into the machine independently of the PilB ATPase^10, 14^ (Fig. 1a). The five pilins interact directly to form a short pilus that is capped by PilY1, which interacts directly with the minor pilins^10^. Pilus extension is initiated by the incorporation of additional major pilin subunits from a reservoir in the IM to the base of the priming complex in a process stimulated by PilB^6, 10, 14^. Conversely, during retraction, major pilin subunits are removed from the base of the pilus and reinserted into the IM in a processs stimulated by PilT^12, 15^. Because the major pilin is added to the priming complex during the initiation of the extension process, the priming complex remains at the tip of the extended pilus^10, 14, 16^. Consistently, PilY1 is involved in surface adhesion, surface-sensing, specificity in host cell recognition during infections, and virulence^14, 16–19^.

**Figure 1.**
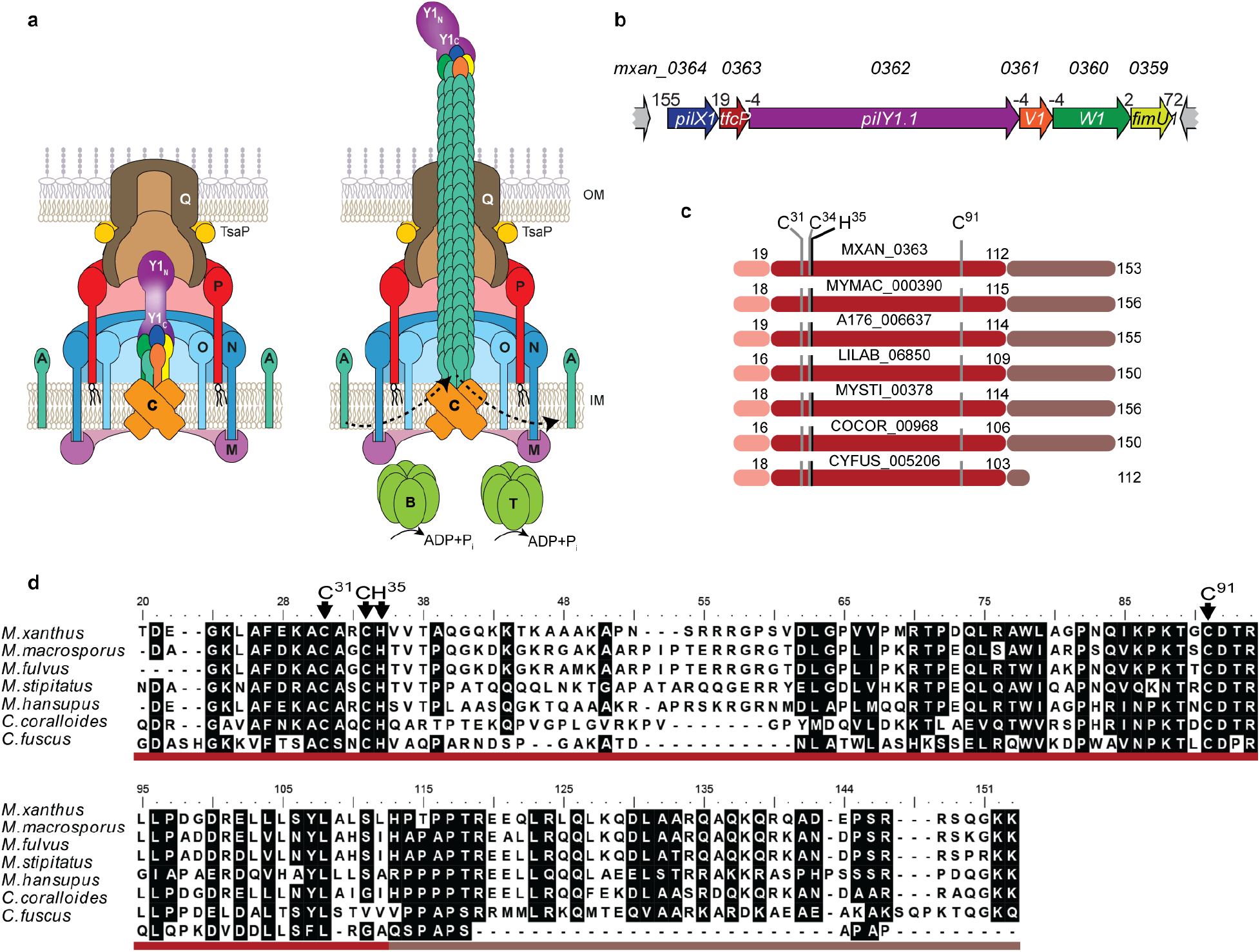
TfcP is a non-canonical cytochrome *c*. **a** Architectural model of non-piliated and piliated T4aPM. PilB and PilT associate with PilC in a mutually exclusive manner during extension and retraction, respectively. Bent arrows, incorporation and removal of the major pilin PilA from the pilus base during extension and retraction, respectively. Proteins labelled with single letters have the Pil prefix. Y1_N_ and Y1_C_ indicate the N- and C-terminal domains of PilY1, respectively. The colour code for the four minor pilins is as in **b**. **b** Genetic organisation of cluster_1 encoding minor pilins, PilY1.1 and TfcP. Locus tags are included above and gene names within genes. Distances between start and stop codons are shown above. **c** Domain architecture of TfcP and homologs. Pink: Type I signal peptide, red: Cytochrome *c* domain, and brown: C-terminal extension. The cytochrome *c* signature motif CxxCH and the distal Cys^91^ residue are indicated. Numbering of amino acids is according to the unprocessed, full-length protein. **d** Sequence alignment of TfcP and homologs. Residues are highlighted based on >80% similarity. Domains are indicated using the color code from **c**. The cytochrome *c* signature motif CxxCH and the distal Cys^91^ residue are indicated. Numbering of amino acids is according to the unprocessed, full-length protein.

Among the 15 proteins of the T4aPM, nine are generally encoded by single copy genes^20^. Some species contain multiple PilT paralogs that enable retractions with different characteristics^21^. The genes for the four minor pilins and PilY1 are also often present in multiple copies^10, 22–24^. The multiplicity of minor pilins and PilY1 proteins has been suggested to allow individual species to assemble priming complexes and tip complexes of different composition and with different properties, thereby allowing the formation of T4aP that can function in a variety of different habitats^10, 14, 25^. Minor pilins are low abundance proteins that share overall structure and sequence homology with the major pilin and have a prepilin signal peptide, a hydrophobic N-terminal α-helix, and a C-terminal globular domain, which is less conserved^26^. PilY1 proteins have a type I signal peptide, are secreted to the periplasm, and are composed of two domains. The conserved C-terminal PilY1-domain adopts a beta-propeller fold^27^ that interacts with the minor pilins in the priming and tip complex^10^ (Fig. 1a). The N-terminal domain is much less conserved and is the domain that mediates host cell recognition, adhesion and surface-sensing^10, 17, 28^.

The soil-dwelling δ-proteobacterium *Myxococcus xanthus* uses T4aP-dependent motility (T4aPdM) and gliding motility to move on surfaces to generate spreading colonies in the presence of nutrients and spore-filled fruiting bodies in the absence of nutrients^29, 30^. The *M. xanthus* genome contains three gene clusters (from here on cluster_1, _2 and _3; proteins labelled with suffix 1, 2 and 3), each encoding four minor pilins and a PilY1 protein^8, 10^. Cluster_1 alone and cluster_3 alone support T4aPdM under standard conditions^10^. While the four respective minor pilins share overall sequence homology, the three PilY1 proteins are highly divergent in their N-terminal domains^10^. Thus, *M. xanthus* has the potential to generate at least two, and possibly three, different T4aPM and T4aP that differ in their priming and tip complexes, respectively.

To understand the functional range of the three minor pilin/PilY1 protein sets, we focused on the proteins of cluster_1. Here, we provide evidence that these proteins form priming and tip complexes in a calcium-dependent manner. We identify the TfcP protein and show that it is a non-canonical cytochrome *c* with an unusual His/Cys heme ligation that is conditionally essential for cluster_1-based T4aP formation and T4aPdM. Specifically, TfcP is important for PilY1.1 stability under low calcium conditions; PilY1.1, in turn, is important for the stability of the cluster_1 minor pilins. The effect of TfcP on PilY1.1 stability depends on calcium binding by PilY1.1 and is bypassed at high calcium concentrations. Our data support a model whereby TfcP is a repurposed cytochrome *c* that promotes calcium binding by PilY1.1 at low calcium concentrations, thereby, allowing cluster_1 to support T4aP function in a broader range of environmental conditions.

## Results

### TfcP is a non-canonical cytochrome *c* important for cluster_1-based T4aP formation

In addition to encoding four minor pilins (PilX1, PilW1, PilV1 and FimU1) and PilY1.1, cluster_1 contains an additional open reading frame (ORF) (Locus tag=*mxan_0363*) (Fig. 1b), for which no homolog is present in cluster_2 and cluster_3. This ORF is conserved in gene clusters encoding minor pilins and PilY1 in other Myxococcales genomes (Fig. S1a). Sequence analysis of Mxan_0363 homologs revealed a type I signal peptide followed by a cytochrome *c* domain that includes a single cytochrome *c* signature motif CxxCH (ref.^31^), and a C-terminal extension enriched in Pro residues and charged amino acids (Fig. 1cd). *C*-type cytochromes are secreted to the periplasm in a Sec-dependent manner where they acquire the heme, which is covalently attached to the two Cys residues in the signature motif by thioether bonds, while the His residue is the proximal axial ligand of the heme iron^32^. ∼90% of cytochromes *c*, the so-called canonical cytochromes *c*, have a Met or His residue ∼60 residues downstream of the signature motif that serves as the second axial ligand of the heme iron^31, 33, 34^. Interestingly, in Mxan_0363 and homologs, this is a Cys residue (Cys^91^ in Mxan_0363) (Fig. 1cd), which is rarely found as the second axial ligand in *c* type cytochromes^34, 35^. In the vicinity of Cys^91^, no conserved Met or His residues are present. All Mxan_0363 homologs except CYFUS_005206 contain the C-terminal extension (Fig. 1cd), which lacks in canonical cytochromes *c*. Thus, Mxan_0363 has features in common with canonical cytochromes *c* but also distinct features. Mxan_0363 homologs were not identified in species other than the listed Myxococcales (Fig. 1cd). From here on, we refer to Mxan_0363 as TfcP for <u>T</u>4aP formation <u>c</u>ytochrome *c* <u>p</u>rotein.

Consistent with the overlap of or short distances between stop and start codons for neighbouring genes (Fig. 1b), fragments were amplified in RT-PCR for all consecutive genes of cluster_1 supporting that they constitute an operon (Fig. S1b).

To test whether TfcP is important for T4aP-formation or function, we generated in-frame deletions of *tfcP* and the remaining five cluster_1 genes. The deletions were generated in a strain in which cluster_2 and cluster_3 had been deleted (Δ2Δ3_cluster strain) because cluster_1 and _3 in the wild-type (WT) strain DK1622 function redundantly to support T4aP formation and T4aPdM^10^. From here on, we used the Δ2Δ3_cluster strain as a reference strain and refer to it as the WT_Δ2Δ3_ strain.

In motility assays for T4aPdM on 0.5% agar supplemented with 0.5% Casitone broth (CTT), WT_Δ2Δ3_ generated the flares at the colony edge characteristic of T4aPdM, while the Δ*pilA* mutant, which lacks the major pilin PilA and served as a negative control, did not (Fig. 2a). As previously shown for cluster_3 genes^10^, T4aPdM was abolished in the Δ*pilX1*, Δ*pilV1*, Δ*pilW1* and Δ*pilY1.1* mutants and reduced in the Δ*fimU1* mutant. Strikingly, T4aPdM was also abolished in the Δ*tfcP* mutant. T4aPdM was restored in all six in-frame deletion mutants by ectopic expression of the relevant gene from a plasmid integrated in a single copy at the Mx8 *attB* site.

**Figure 2.**
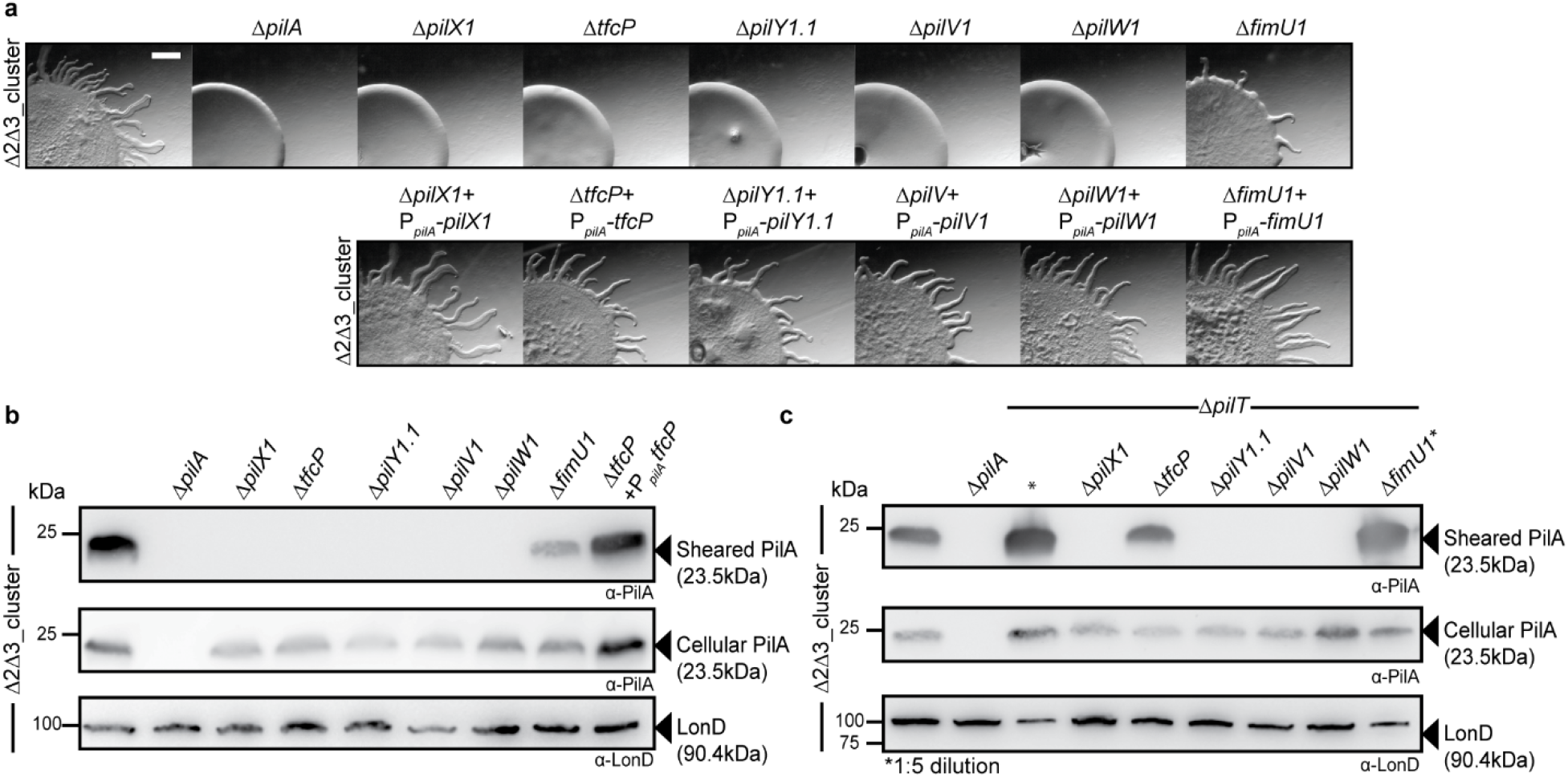
TfcP, minor pilins and PilY1.1 of cluster_1 are important for T4aPdM and T4aP formation. **a** Assay for T4aPdM. WT_Δ2Δ3_ and strains with deletions of individual cluster_1 genes, and the corresponding complementation strains were spotted on 0.5% agar supplemented with 0.5% CTT and imaged after 24 hrs. Scale bar, 1 mm. **b** Shearing assay for T4aP formation. T4aP sheared off from ∼15 mg cells grown on 1.5% agar supplemented with 1.0% CTT were separated by SDS-PAGE and probed with α-PilA antibodies (top panel). Middle panel, 40 µg of protein from total cell extracts separated by SDS-PAGE and probed with α-PilA antibodies and, after stripping, with α-LonD antibodies as a loading control (lower panel). **c** Shearing assay for T4aP formation in retraction deficient strains. T4aP-formation was assayed as in **b**. In lanes labeled with *, five-fold less protein was loaded.

To pinpoint the mechanism causing the T4aPdM defect in the cluster_1 mutants, we assessed T4aP formation in the six in-frame deletion mutants using a shearing assay. In this assay, T4aP are sheared off the cell surface, and the level of the major pilin PilA in the sheared fraction quantified by immuno-blot analysis (Fig. 2b). None of the five non-motile mutants formed detectable T4aP, while the Δ*fimU1* mutant still assembled T4aP at a much-reduced level compared to the parent strain. For all six in-frame deletion mutants, the total cellular level of PilA was similar or slightly lower than in the parent WT_Δ2Δ3_ strain. As expected, T4aP-formation in the Δ*tfcP* mutant was complemented by ectopic expression of *tfcP*.

To distinguish whether the defect in T4aP-formation was caused by lack of extension or by hyper-retractions, we examined T4aP-formation in the six in-frame deletion mutants additionally containing a Δ*pilT* mutation and, thus, lacking the PilT retraction ATPase (Fig. 2c). The WT_Δ2Δ3_Δ*pilT* strain formed T4aP at a highly increased level compared to WT_Δ2Δ3_ consistent with previous observations for the WTΔ*pilT* strain^12^. In the absence of PilT, T4aP-formation was partially restored in the Δ*tfcP* mutant, but at a much-reduced level compared to the WT_Δ2Δ3_Δ*pilT* strain. By contrast, T4aP-formation in the Δ*pilX1*, Δ*pilV1*, Δ*pilW1 and* Δ*pilY1.1* mutants was not restored. For all in-frame deletion mutants except for the Δ*fimU1* mutant, the total cellular level of PilA was lower than in WT_Δ2Δ3_Δ*pilT* strain. We conclude that TfcP is important but not essential for cluster_1-dependent T4aP extension while the three minor pilins PilX1, -V1 and - W1 as well as PilY1.1 are essential for T4aP-formation, and FimU1 plays a less important role. The observations are in agreement with similar experiments involving minor pilins and PilY1.3 of cluster_3^10^.

### TfcP is important for PilY1.1 stability

To understand how TfcP might be involved in T4aP extension, we used proteomics on whole-cell extracts to quantify the accumulation of T4aPM components in WT_Δ2Δ3_ and WT_Δ2Δ3_Δ*tfcP* strains. To increase sensitivity, we used targeted proteomics in which protein abundance is quantified relative to heavy labelled reference peptides of the proteins of interest (Methods). In absence of TfcP, accumulation of 10 T4aPM components was largely unaffected, while the accumulation of the four minor pilins and PilY1.1 was strongly reduced (Fig. 3a). Because PilY1 of cluster_3 is important for the stability of cluster_3 minor pilins^10^, we performed targeted proteomics on the WT_Δ2Δ3_Δ*pilY1.1* strain. In this strain, accumulation of the four minor pilins was also strongly reduced, while TfcP accumulation was increased (Fig. 3a). In immuno-blot analysis, we observed that in the absence of individual cluster_1 minor pilins, accumulation of TfcP was increased and PilY1.1 unchanged (Fig. 3b). Immuno-blot analysis also confirmed that PilY1.1 accumulation was strongly reduced in the absence of TfcP while TfcP accumulation was increased in the absence of PilY1.1 (Fig. 3b). Altogether, these observations support that TfcP is important for accumulation of PilY1.1, which, in turn, is important for minor pilin accumulation.

**Figure 3.**
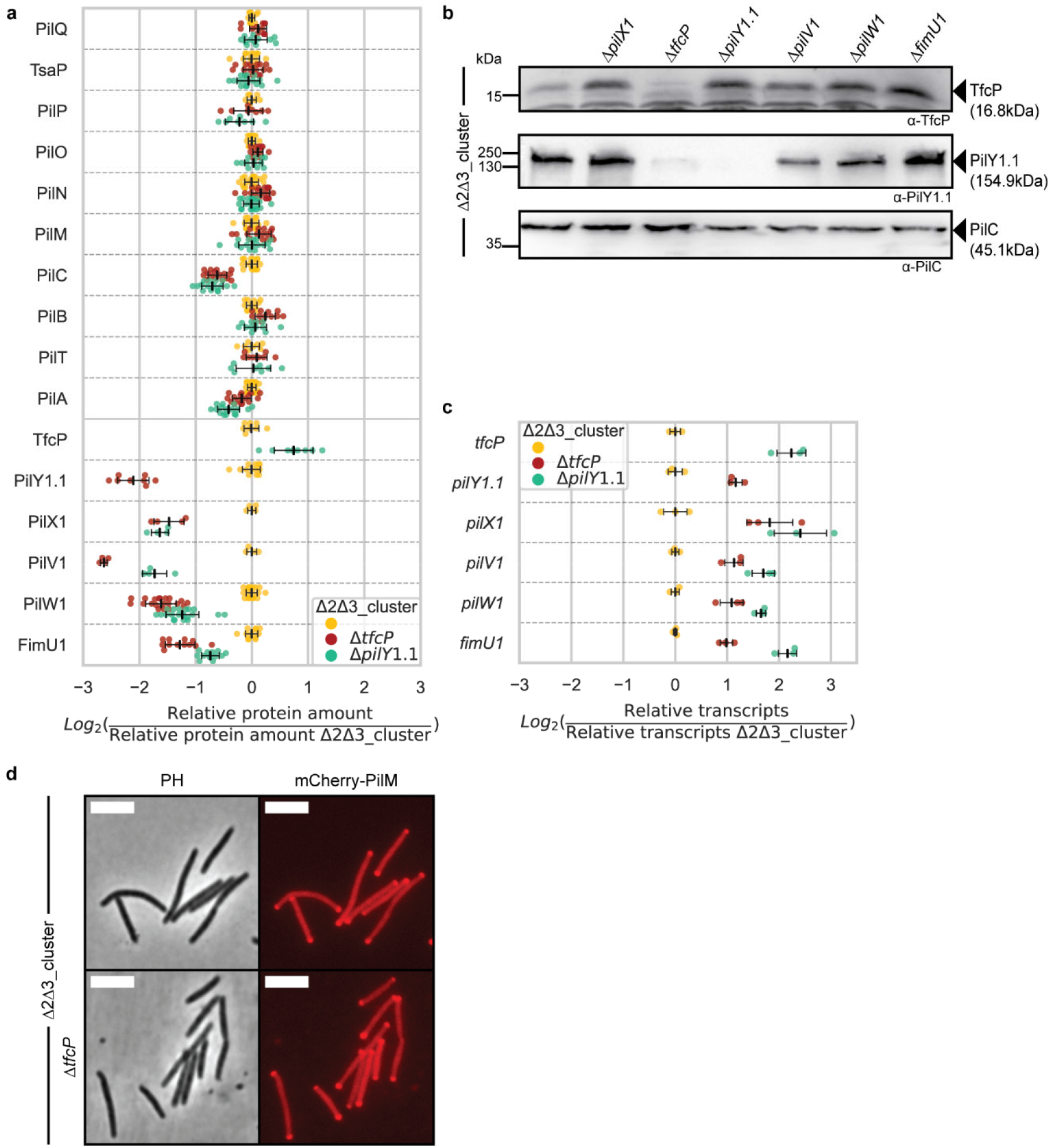
TfcP is important for stability of PilY1.1 and minor pilins of cluster_1. **a** Accumulation of proteins of the T4aPM and cluster_1. Cells were grown in 1.0% CTT suspension culture. Relative protein amounts were determined using targeted proteomics with one to five heavy labelled reference peptides for each protein spiked into the trypsin-digested cell lysates (Methods). To calculate relative protein amounts, the light-to-heavy intensity ratio of the endogenous (light) and reference (heavy) peptide was calculated. For every strain, four biological replicates were analysed. Individual data points represent the log_2_ ratio of the relative amount of one peptide in one biological replicate to the mean relative amount of the same peptide in the WT_Δ2Δ3_ strain (calculated from the four biological replicates). Center marker and error bars in black: Mean and standard deviation (STDEV) of all values for one protein. **b** Immuno-blot analysis of TfcP and PilY1.1 accumulation. Cells were grown in 1.0% CTT suspension culture. Total cell extracts from the same number of cells were separated by SDS-PAGE and analysed by immuno-blotting. PilC was used as loading control. **c** qRT-PCR analysis of transcript levels of cluster_1 genes. Total RNA was isolated from cells grown in 1.0% CTT suspension culture. Individual data points represent three biological replicates with each two technical replicates, and in which the ratio of the relative transcript level in a mutant over the transcript level in the WT_Δ2Δ3_ strain is plotted. Center marker and error bars: Mean and STDEV. **d** Localisation of mCherry-PilM in the Δ*tfcP* strain. Cells were grown in 1.0% CTT suspension culture, placed on 1.0% agarose supplemented with TPM, and immediately imaged by phase contrast (PH) and fluorescence microscopy. Scale bar, 5 µm.

To resolve whether the effect of the Δ*tfcP* mutation on PilY1.1 and the Δ*pilY1.1* mutation on minor pilin accumulation was due to altered transcription of the relevant genes or altered protein stability, we performed qRT-PCR analysis on total RNA from the WT_Δ2Δ3_, WT_Δ2Δ3_Δ*tfcP* and WT_Δ2Δ3_Δ*pilY1.1* strains. Transcript levels of the cluster_1 genes were increased in the Δ*tfcP* and the Δ*pilY1.1* mutants (Fig. 3c), suggesting negative feedback regulation of cluster_1 genes. While the mechanism involved in this regulation remains unresolved, these results do not support that the reduced levels of PilY1.1/minor pilins and minor pilins in the absence of TfcP and PilY1.1, respectively are caused by reduced synthesis. Rather they support that TfcP stabilises PilY.1.1, which, in turn, stabilises the four minor pilins. Accumulation dependencies have also been reported for the cluster_3 proteins in which PilY1.3 and minor pilins interact directly to mutually stabilise each other^10^.

In *M. xanthus*, the T4aPM assembles at the two poles^10, 36–38^. To exclude that deletion of *tfcP* affects assembly of the T4aPM, we used the bipolar localisation of the cytoplasmic protein PilM as proxy for T4aPM assembly^37^. We observed bipolar localisation of an active mCherrry-PilM fusion in most cells of the WT_Δ2Δ3_ and WT_Δ2Δ3_Δ*tfcP* strains (Fig. 3d), supporting that TfcP is not important for assembly of the remaining proteins into rudimentary T4aPM.

### TfcP is a periplasmic protein

To understand how TfcP stabilises PilY1.1, we determined its subcellular localisation using active TfcP-FLAG and TfcP-sfGFP fusions expressed at native or slightly above native levels from the endogenous locus (Fig. 4ab). Cells of WT_Δ2Δ3_ synthesizing TfcP-FLAG were fractionated into fractions enriched for soluble, IM and OM proteins. TfcP-FLAG was present in the soluble fraction while the control proteins fractionated as described^12, 36^ (Fig. 4c). Because sequence analysis of TfcP predicted a type I signal peptide, this experiment supports that TfcP localises to the periplasm similarly to PilY1 proteins^10^. In agreement with these observations, in fluorescence microscopy, TfcP-sfGFP localised along the entire cell periphery and foci formation was not observed (Fig. 4d).

**Figure 4.**
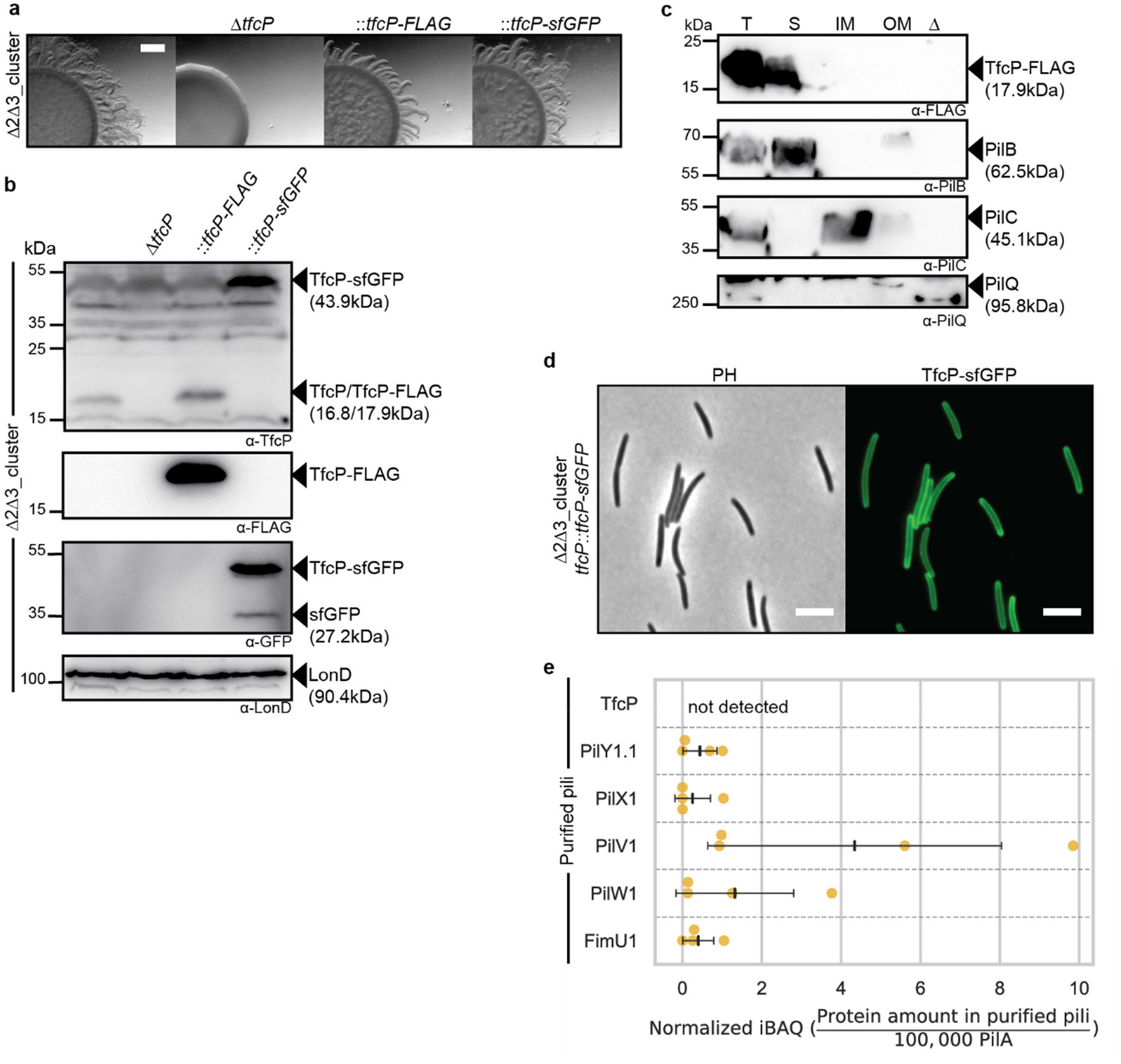
TfcP is a periplasmic protein. **a** Assay for T4aPdM. Strains were assayed as in Fig. 2a. Scale bar, 1 mm. **b** Immuno-blot analysis of TfcP-FLAG and TfcP-sfGFP accumulation. Cells were grown in 1.0% CTT suspension culture and analysed as in Fig. 3b. **c** Subcellular localisation of TfcP-FLAG. Cells were grown in 1.0% CTT suspension culture and fractionated into fractions enriched for soluble proteins (S), IM proteins (IM) and OM proteins (OM). T indicates total cell extract. In the lane marked Δ, total cell extract of the Δ*pilBTCMNOPQ* mutant was used as negative control. Protein from the same number of cells was loaded per lane, and analysed by immuno-blotting. PilB, PilC and PilQ serve as controls for the fractionation and localise to the cytoplasm, IM and OM, respectively^12, 36^. **d** Determination of TfcP-sfGFP localisation. Cells were grown in 1.0% CTT suspension culture, and analysed as in Fig. 3d. Scale bar, 5 µm. **e** Label-free quantification (LFQ) of cluster_1 proteins in purified pili. Pili were isolated from cells grown on 1.5% agar supplemented with 1.0% CTT. Normalised iBAQ (intensity based absolute quantification) values (Methods) were determined in four biological replicates for WT_Δ2Δ3_Δ*pilT* and the negative control WT_Δ2Δ3_Δ*pilT*Δ*pilB*. iBAQ values of WT_Δ2Δ3_Δ*pilT* were background corrected by subtraction of the mean iBAQ value of the four replicates of the negative control and rescaled to the iBAQ value of 100,000 PilA molecules in the same sample. Center marker and error bars: Mean and STDEV.

To determine whether TfcP is present in pili, we purified pili from the WT_Δ2Δ3_Δ*pilT* mutant (Fig. S2a) and used label-free quantitative proteomics to quantify cluster_1 proteins. All cluster_1 proteins except TfcP were detected in low amounts relative to the PilA major pilin (Fig. 4e) as described for cluster_3 proteins^10^.

Altogether, these observations support that the minor pilins and PilY1.1 of cluster_1 form a priming complex in the T4aPM for T4aP extension as well as a pilus tip complex. We also conclude that TfcP is a soluble, periplasmic protein that stabilises PilY1.1, and that TfcP is incorporated into neither the T4aPM nor T4aP. These findings together with previous results that all proteins that are incorporated into the T4aPM localise (bi)polarly^10, 36–38^, suggest that the stabilising effect of TfcP on PilY1.1 occurs in the periplasm and before PilY1.1 incorporation into the T4aPM.

### TfcP is a non-canonical cytochrome *c* with a low redox potential and heme binding is important for TfcP stability *in vivo*

We overexpressed and purified a MalE-tagged TfcP variant (MalE-TfcP) from *Escherichia coli* to assess its heme-binding and redox characteristics (Fig. S2b). The fusion for overexpression contains the MalE type I signal peptide and lacks the TfcP signal peptide. Purified MalE-TfcP exhibited a distinct red colour indicating that it binds heme (Fig 5a). Oxidised MalE-TfcP had strong peroxidase activity (Fig. 5a) in agreement with heme-containing proteins having intrinsic peroxidase activity^39^. Importantly, peroxidase activity was inhibited when MalE-TfcP was reduced by dithiothreitol (DTT), supporting that this activity results from oxidised heme bound to MalE-TfcP^40^.

**Figure 5.**
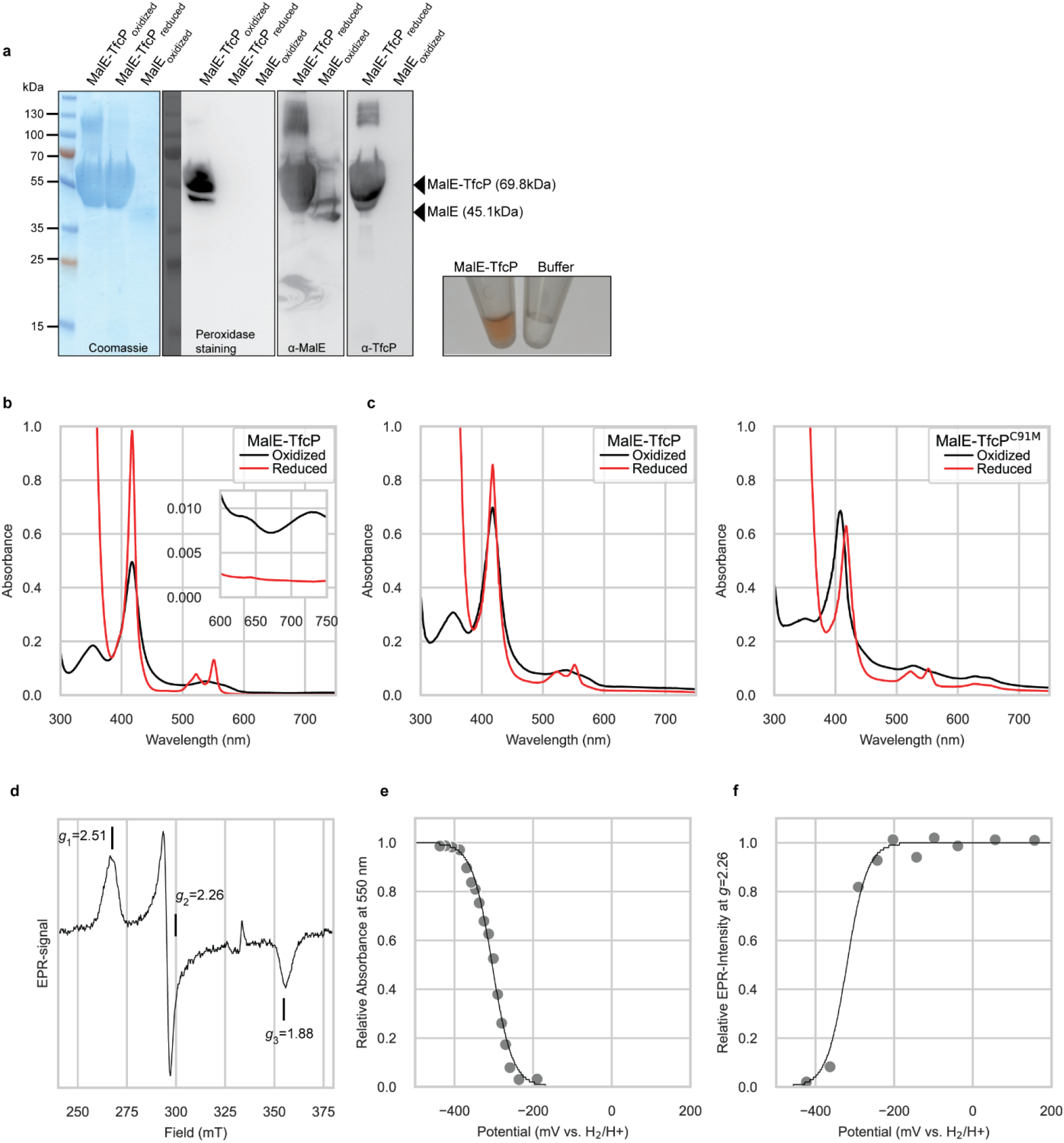
TfcP is a redox active, heme-binding protein. **a** TfcP heme-binding assay. Panels from left-to-right, MalE-TfcP in oxidised (as purified) and reduced state (after addition of DTT) stained with Coomassie Blue, analysed for heme-binding by peroxidase staining using a luminescent horse radish peroxidase (HRP) substrate and MalE as negative control, detected by immuno-blotting with α-MalE and α-TfcP as indicated, and image of purified MalE-TfcP in buffer. **b** UV-Vis spectra of purified MalE-TfcP in the oxidised and reduced (after addition of sodium-dithionite) state. Inset shows the absorbance in the 600-750 nm region. Experiment was done using a Shimadzu 1900 spectrophotometer. **c** UV-Vis spectra of purified MalE-TfcP variants in the oxidised and reduced state. Experiments were done using a Tecan200Pro platereader and, therefore, the spectrum of WT TfcP is included again. **d** EPR spectrum of MalE-TfcP. Spectra were recorded in the oxidised state at 12 K, 0.32 mW microwave power, 1.5 mT modulation amplitude (9.3523 GHz). **e** Redox titration of MalE-TfcP. The 550 nm absorbance at 23°C is plotted versus the solution potential and fitted to the Nernst equation. **f** Redox titration of MalE-TfcP following the EPR-intensity at *g*=2.26 of samples poised at indicated solution redox potentials.

To assess the heme-binding properties of TfcP, we used UV-Vis spectroscopy. MalE-TfcP has a cytochrome *c*-like spectrum with a strong Soret-peak in the oxidised form and after reduction with sodium-dithionite (Fig. 5b). In the spectrum of reduced TfcP, the α- and β-peak become visible in the 550 nm region. This fits well to spectra of canonical cytochromes *c*. Interestingly, we did not observe a red-shift of the Soret-peak from the oxidised to the reduced spectrum. For canonical cytochromes *c* with His/His or His/Met coordination a 10 nm bathochromic shift is typically observed, while a semisynthetic cytochrome *c* with His/Cys coordination of the heme iron did not exhibit this shift^35, 41, 42^, suggesting that Cys^91^ (Fig. 1cd) is the second axial ligand in TfcP and responsible for the lack of the red-shift. This is also supported by the presence of a peak at 360 nm in the oxidised spectrum, which has been reported for His/Cys ligation^43^. The presence of cysteine-to-Fe^3+^ charge transfer bands at ∼630 nm and ∼730 nm in the oxidised form, which disappear upon dithionite reduction (Fig. 5b, inset), are also in full agreement with spectral properties of His/Cys coordinated *c*-type cytochromes^35^. To further support that the special spectral properties are due to Cys^91^, we purified the MalE-TfcP^C91M^ variant (Fig. S2b). In this variant, a red-shift was observed upon reduction of the protein (Fig 5c). In addition, the 360 nm peak was not detected in the oxidised form. We conclude that Cys^91^ is the second axial ligand of the heme iron in TfcP and that TfcP is an unusual cytochrome *c*.

We used electron paramagnetic resonance (EPR)-spectroscopy to investigate the environment of the heme-center. We obtained *g*-values of 2.51, 2.26 and 1.88 (Fig. 5d) that fit well to the *g*-values observed for multiple heme-containing proteins with a cysteine thiolate ligated heme iron^44^. Cytochromes *c* with a distal Cys were reported to have a very low midpoint potential in the range of −350 mV while canonical cytochromes *c* have a potential of approximately +250 mV^35, 41, 42, 45^. To determine whether TfcP has a similarly low redox-potential, we used UV-Vis and EPR monitored redox titrations. For the UV-Vis redox-titration, MalE-TfcP was incubated with a redox-mediator cocktail. Spectra and potentials were recorded in five-minute intervals after addition of sodium-dithionite. After plotting the absorbance change at 550 nm versus the potential and fitting to the Nernst equation, the midpoint potential was determined as E_m_ = −304±8 mV, where 8 mV represent the intrinsic fitting error in one experiment (Fig. 5e). In the EPR-monitored redox titration, we followed the change of *g*=2.26 EPR signal in frozen samples obtained by sequential reduction with sodium-dithionite in the presence of mediators and found a midpoint potential of E_m_ = −320±15 mV, where 15 mV represent the intrinsic fitting error in one experiment. Overall, both experiments support that TfcP has a very low redox-potential. The slight difference between the two experiments is likely due to pH changes, which occur during freezing. The low redox potential (approx. −312 mV) indicates that TfcP is not likely to be part of a respiratory chain in *M. xanthus* (see Discussion).

To clarify whether the heme-binding characteristics of TfcP is important for activity *in vivo*, we substituted the two Cys residues to Ala in the C^31^xxCH heme-binding motif and Cys^91^ to His or Met (Fig. 1cd). The variants were synthesised ectopically as FLAG-tagged proteins in the Δ*tfcP* mutant from the strong *pilA* promoter. None of the variants complemented the motility defect in the Δ*tfcP* mutant, while the WT protein did (Fig. S3a). However, all three mutant variants accumulated at much-reduced levels compared to TfcP and TfcP-FLAG expressed from the native site, and none of the three mutant variants supported PilY1.1 accumulation (Fig. S3b). These observations demonstrate that heme-binding and distal coordination of the heme iron are important for TfcP stability.

Also, a FLAG-tagged TfcP^Δ118–153^ variant lacking the C-terminal extension was not able to complement the T4aPdM defect in the Δ*tfcP* mutant (Fig. S3a), accumulated at a reduced level, and did not support PilY1.1 accumulation (Fig. S3b), providing experimental support for the functional relevance of this C-terminal extension.

### Added calcium restores T4aP-formation in the absence of TfcP

Several PilY1 proteins have been shown to bind calcium using an EF-hand-like motif in the C-terminal domain, and calcium binding is important for function^27, 46, 47^. PilY1.1 and PilY1.2 also contain the consensus EF-hand-like calcium binding Dx[DN]xDGxxD motif in the C-terminal PilY1 domain, while PilY1.3 has two calcium binding motifs in the N-terminal domain (Fig. 6a). In a homology model of the C-terminal domain of PilY1.1, the D^1165^xDxDNxxD^1173^ motif is located in a surface exposed loop between two β–strands as described for the PilY1 domain of *Pseudomonas aeruginosa*^27^ (Fig. 6b).

**Figure 6.**
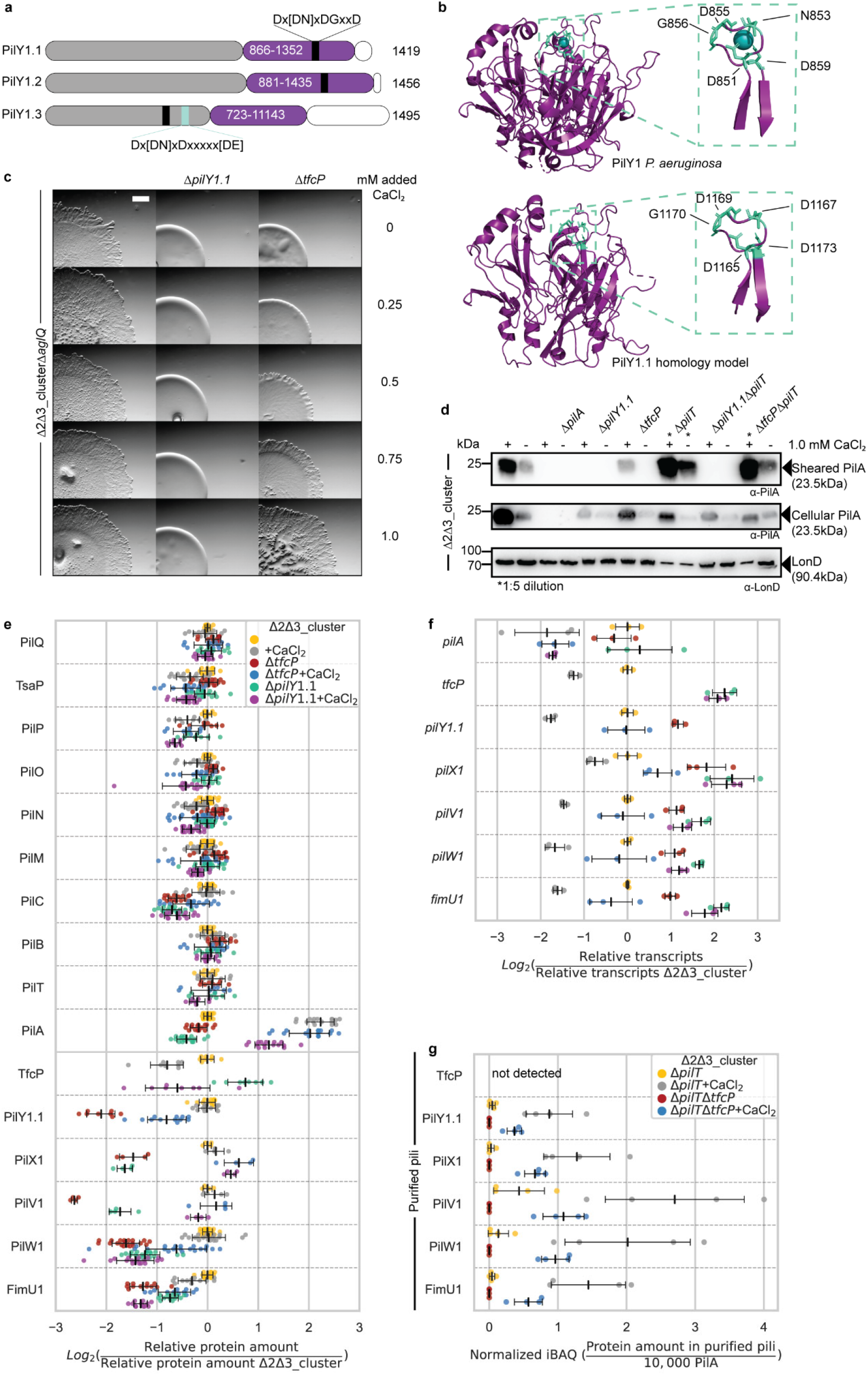
Added CaCl_2_ compensates for lack of TfcP. **a** Domain architecture of PilY1 proteins of *M. xanthus*. Purple, C-terminal PilY1 domain; grey, N-terminal domains; white; C-terminal sequences. EF hand-like calcium binding motif is in black together with the consensus sequence; light blue box indicates the second calcium binding motif in PilY1.3 together with the consensus sequence^25^. **b** Comparison of PilY1 structure of *P. aeruginosa* (PDB: 3HX6)^27^ and a homology model of PilY1.1; inset, zoom of calcium binding motif. **c** Assay for T4aPdM. Cells were grown in 1.0% CTT suspension culture and plated on 0.5% agar supplemented with 0.5% CTT without or with added CaCl_2_ and imaged after 24 hrs. Final concentrations of added CaCl_2_ are indicated. Scale bar, 1 mm. **d** Shearing assay for T4aP formation. T4aP sheared off from ∼15 mg cells grown on 1.5% agar supplemented with 1.0% CTT and 1.0 mM CaCl_2_ as indicated, and analysed as in Fig. 2b. **e** Accumulation of proteins of the T4aPM. Cells were grown in 1.0% CTT suspension culture without or with 1.0 mM added calcium as indicated. Proteins were quantified as in Fig. 3a. Data for samples without added CaCl_2_ are the same as in Fig. 3a and included for comparison. **f** qRT-PCR analysis of transcript levels of cluster_1 genes and *pilA*. Total RNA was isolated from cells grown in 1.0% CTT suspension culture without or with added calcium as indicated. Transcripts were quantified as in Fig. 3c. Colour code is as in **e**. Data for samples without added CaCl_2_ are the same as in Fig. 3e and included for comparison. **g** LFQ proteomics of cluster_1 proteins in purified pili. Pili were isolated from cells grown on 1.5% agar supplemented with 1.0% CTT without or with added CaCl_2_ as indicated. Normalised iBAQ values were calculated as in Fig. 4e and background corrected by subtraction of the mean iBAQ value of the four replicates of the relevant negative control, and rescaled to 10,000 PilA molecules in the same sample. Data for WT_Δ2Δ3_ without added CaCl_2_ are the same as in Fig. 4e and included for comparison.

To address the effect of calcium on T4aPdM, we first considered that the previous experiments were performed either in 1.0% CTT (targeted proteome analyses and qRT-PCR), which has a calcium concentration of ∼30 µM according to the manufacturer, or on 0.5% agar supplemented with 0.5% CTT (motility assays) or 1.5% agar supplemented with 1.0% CTT (T4aP purification). The estimated calcium concentration of 0.5% agar is ∼0.15 mM^48^. Next, we repeated the assays for T4aPM in the presence of additional CaCl_2_. To ensure that the effect of added CaCl_2_ could be attributed to T4aP, we used the WT_Δ2Δ3_Δ*aglQ* strain, which lacks the AglQ motor for gliding^49, 50^. In the presence of ≥0.25 mM added CaCl_2_, WT_Δ2Δ3_Δ*aglQ* exhibited a dramatic change of motility pattern from expansion in flares to a radial film-like expansion (Fig. 6c). Intriguingly, the WT_Δ2Δ3_Δ*aglQ*Δ*tfcP* mutant also responded to increasing calcium concentrations. At added CaCl_2_ concentrations ≥0.5 mM, this mutant regained T4aPdM and at 1.0 mM displayed a motility pattern similar to that of the parent strain. By contrast, added CaCl_2_ did not restore T4aPdM in the WT_Δ2Δ3_Δ*pilY1.1* mutant even at 10 mM (Fig. 6c; Fig. S4ab). Likewise, 10 mM CaCl_2_ did not restore T4aPdM in the Δ*pilX1*, Δ*pilV1*, Δ*pilW1 and* Δ*pilA* mutants while the Δ*fimU1* mutant displayed the same radial motility pattern as the parent strain (Fig. S4a). In additional experiments, we observed that WT_Δ2Δ3_Δ*aglQ* responded to added CaCl_2_ concentrations as low as 0.025 mM while the WT_Δ2Δ3_Δ*aglQ*Δ*tfcP* only responded at ≥0.5 mM (Fig. S4b). In control experiments, we observed that neither 5 mM NaCl nor 5 mM MgCl_2_ restored motility in the WT_Δ2Δ3_Δ*tfcP* mutant. We also observed that a strain only containing cluster_3 responded to added CaCl_2_ with an altered expansion pattern; however, this pattern was only evident at added CaCl_2_ concentrations ≥0.25 mM (Fig. S4b). We conclude that CaCl_2_ at an added concentration of 1.0 mM restores T4aPdM in the WT_Δ2Δ3_Δ*tfcP* strain. From here on, we used an added CaCl_2_ concentration of 1.0 mM.

Consistent with the effect of added CaCl_2_ on motility, the WT_Δ2Δ3_Δ*tfcP* mutant formed T4aP in the presence of 1.0 mM CaCl_2_ based on the shearing assay, while the WT_Δ2Δ3_Δ*pilY1.1* mutant did not (Fig. 6d). Added CaCl_2_ also increased the amount of shearable pili in WT_Δ2Δ3_. The level of T4aP in the WT_Δ2Δ3_Δ*tfcP* mutant was lower than in WT_Δ2Δ3_ (Fig. 6d). Calcium also increased the amount of total cellular PilA in all strains (Fig. 6d). The retraction deficient strains WT_Δ2Δ3_Δ*pilT* and WT_Δ2Δ3_Δ*tfcP*Δ*pilT* assembled more T4aP in the presence of added CaCl_2_ than in the absence supporting that calcium stimulates T4aP-formation rather than counteracting retractions (Fig. 6d). We conclude that 1.0 mM added CaCl_2_ can substitute for TfcP function in T4aP-formation and T4aPdM.

### TfcP enhances calcium-dependent stabilisation of PilY1.1

To understand how a high concentration of calcium compensates for lack of TfcP, we used targeted proteomics. In WT_Δ2Δ3_, 1.0 mM CaCl_2_ caused an increase in PilA abundance and a decrease in TfcP abundance while accumulation of other T4aPM components including the remaining cluster_1 proteins was largely unaffected (Fig. 6e). In the WT_Δ2Δ3_Δ*tfcP* mutant, added CaCl_2_ not only caused an increase in PilA abundance but also increased the abundance of all remaining cluster_1 proteins including PilY1.1 (Fig. 6e). In the WT_Δ2Δ3_Δ*pilY1.1* mutant, extra CaCl_2_ also caused increased PilA abundance, but a reduction in TfcP abundance as in the WT_Δ2Δ3_ parent strain. PilW1 and FimU1 abundance was unaffected by added CaCl_2_ in WT_Δ2Δ3_Δ*pilY1.1*, while PilV1 and PilX1 abundance increased. We conclude that a high concentration of CaCl_2_ causes increased PilY1.1 accumulation in the absence of TfcP. In addition, extra CaCl_2_ also caused (1) increased PilA accumulation independently of TfcP and PilY1.1, (2) decreased accumulation of TfcP independently of PilY1.1, and (3) increased accumulation of the minor pilins PilX1 and PilV1 independently of PilY1.1.

Because changes in extracellular calcium can cause altered gene expression^51^, we performed qRT-PCR analyses to discriminate whether added CaCl_2_ affects transcription or protein stability. We observed significant changes in the transcription of all cluster_1 genes as well as of *pilA* in response to added CaCl_2_ (Fig. 6f); however, generally, these changes did not correlate with the altered protein accumulation profiles. For instance, in WT_Δ2Δ3_, 1.0 mM added CaCl_2_ caused (1) increased PilA accumulation but *pilA* transcription was decreased, and (2) decreased *pilY1.1* transcription but PilY1.1 abundance remained unchanged, and in the WT_Δ2Δ3_Δ*tfcP* mutant, added CaCl_2_ caused decreased *pilY1.1* transcription but PilY1.1 abundance increased. We conclude that added CaCl_2_ at 1.0 mM can substitute for TfcP in stabilising PilY1.1.

Label-free quantitative proteomics of purified pili from WT_Δ2Δ3_Δ*pilT* and WT_Δ2Δ3_Δ*pilT*Δ*tfcP* (Fig. S2a) revealed a strong increase in the abundance of cluster_1 minor pilins and PilY1.1 relative to PilA in the presence of calcium (Fig. 6g) suggesting that calcium also stabilises minor pilins and PilY1.1 in the tip complex. Of note, TfcP was also not detected in purified pili from WT_Δ2Δ3_Δ*pilT* grown in the presence of added calcium (Fig. 6g).

To determine whether the effect of calcium on PilY1.1 stability depends on its binding to PilY1.1, we attempted to purify full-length PilY1.1 or its C-terminal domain but were unsuccessful, thus, precluding *in vitro* analyses of the calcium binding properties of PilY1.1. Therefore, to assess calcium binding by PilY1.1 *in vivo*, we introduced the Asp^1173^ to Ala substitution in the EF-hand-like calcium binding D^1165^xDxDNxxD^1173^ motif in PilY1.1 (Fig. 6b) and expressed the protein from the native site in WT_Δ2Δ3_ strains. The homologous substitution in other PilY1 proteins disrupts calcium binding without affecting the overall structure of the C-terminal beta-propeller _domain27,46,47._

The *pilY1.1*^D1173A^*tfcP*^+^ mutant was strongly reduced in T4aPdM in the absence of added CaCl_2_ (Fig. 7a); however, this strain regained T4aPdM and was indistinguishable from the parent strain at ≥0.25 mM added CaCl_2_. By contrast, the *pilY1.1*^D1173A^Δ*tfcP* strain was non-motile even at 10 mM added CaCl_2_. These observations support that calcium binding is important for PilY1.1 function and that PilY1.1^D1173A^ is fully functional at elevated calcium concentrations but only if TfcP is present.

**Figure 7.**
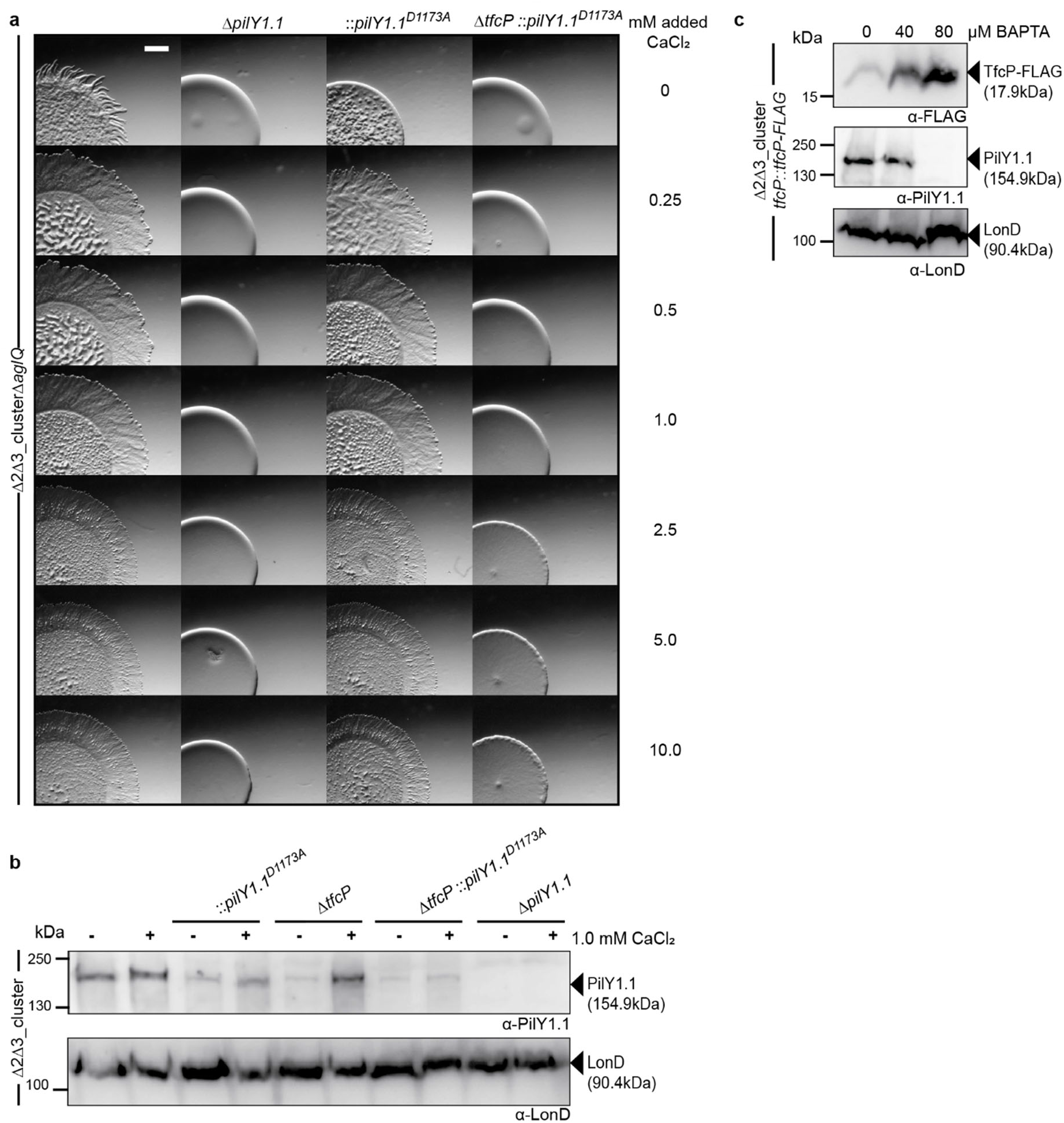
Calcium binding by PilY1.1 is essential for TfcP function. **a** Assay for T4aPdM. Cells were grown in 1.0% CTT suspension culture and plated on 0.5% agar supplemented with 0.5% CTT and imaged after 24 hrs. The final concentration of added CaCl_2_ is indicated. Scale bar, 1 mm. **b** Accumulation of PilY1.1 variants. Cells were grown in 1.0% CTT suspension culture without or with 1.0 mM CaCl_2_, total cell extract isolated and analysed by immuno-blot as in Fig. 3b. **c** Accumulation of TfcP-FLAG and PilY1.1 in presence of BAPTA. Cells were grown in 1.0% CTT in suspension, exposed to indicated concentrations of BAPTA for 16 hrs, total cell extract isolated, and analysed by immuno-blot as in Fig. 3b.

Consistent with the observations for T4aPdM, PilY1.1^D1173A^ accumulation was reduced in the *pilY1.1*^D1173A^*tfcP*^+^ mutant in the absence of added CaCl_2_, and 1.0 mM CaCl_2_ at least partially restored its accumulation (Fig. 7b). By contrast, in the *pilY1.1*^D1173A^Δ*tfcP* strain, PilY1.1^D1173A^ was detected at very low levels in the absence of added CaCl_2_ and did not increase upon addition of CaCl_2_. Thus, PilY1.1^D1173A^ depends on TfcP for stability, and responds to added calcium only in the presence of TfcP. By comparison, PilY1.1^WT^, is fully functional at ≥1.0 mM added CaCl_2_ in the absence of TfcP (Fig. 6c).

Finally, to determine whether TfcP can stabilise PilY1.1^WT^ independently of calcium, we analysed PilY1.1 accumulation in the presence of the highly specific calcium chelator BAPTA (1,2-bis(o-aminophenoxy)ethane-N,N,N′,N′-tetraacetic acid). In WT_Δ2Δ3_ cells expressing TfcP-FLAG from the endogenous site and grown in 1.0% CTT, PilY1.1 was still detected in the presence of 40 µM but not in the presence of 80 µM BAPTA while TfcP was detected under all conditions and increased upon BAPTA addition (Fig. 7c). These observations strongly support that TfcP can only stabilise PilY1.1 in the presence of calcium. Because CaCl_2_ can stabilise PilY1.1 in the absence of TfcP, these observations suggest that the primary function of TfcP is to chaperone calcium binding by PilY1.1 at low calcium concentrations.

## Discussion

Here, we identify TfcP, a repurposed, non-canonical cytochrome *c*, as a novel protein important for cluster_1-dependent T4aP formation in *M. xanthus* at low calcium concentrations. We demonstrate that TfcP stabilises PilY1.1 at low calcium concentrations. PilY1.1, in turn, stabilises the four minor pilins of cluster_1 in that way enabling the formation of the cluster_1-based priming complex in the T4aPM and, thus, T4aP formation. Bacteria in their natural habitats experience large fluctuations in environmental conditions and depend on adaptive strategies to endure such changes. TfcP expands the range of calcium concentrations under which cluster_1 encoded minor pilins and PilY1.1 can support T4aPdM thereby increasing fitness of *M. xanthus* under changing environmental conditions and enabling colonisation of habitats with low calcium concentrations (Fig. 8).

**Figure 8.**
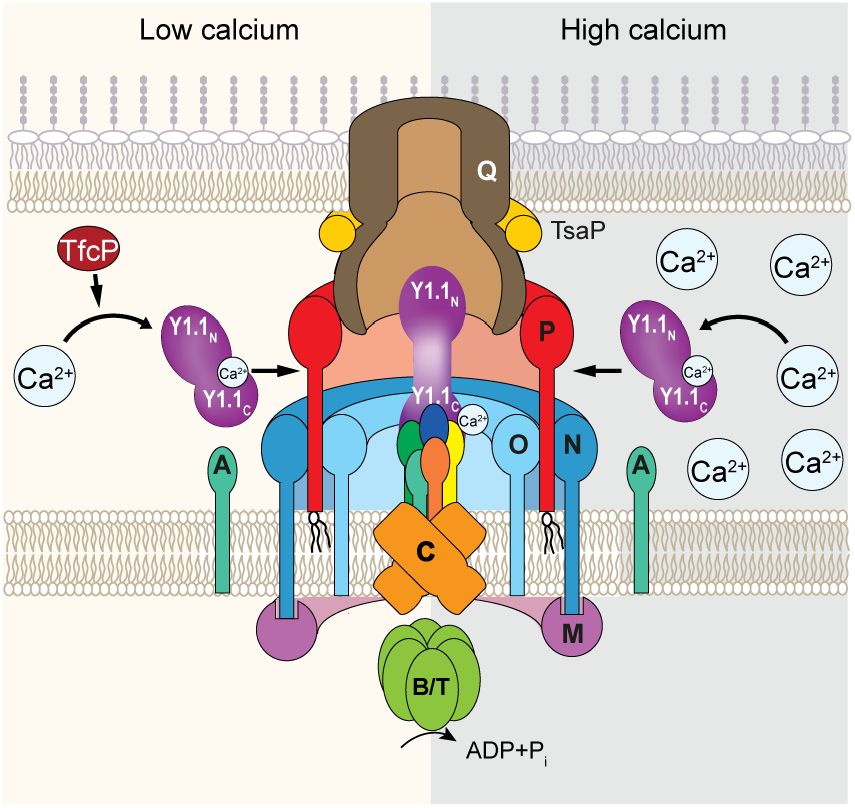
Model of TfcP function at low and high calcium concentrations. For simplicity, PilB and PilT are not shown separately. Y1.1_N_ and Y1.1_C_ indicate the N- and C-terminal domains of PilY1, respectively. The colour code for the four minor pilins is as in Fig. 1b.

Several lines of evidence support that that the effect of TfcP on PilY1.1 stability is calcium-dependent. First, under standard conditions, *M. xanthus* is exposed to ∼30 µM calcium in CTT suspension culture and ∼0.15 mM calcium on 0.5% agar-plates for motility assays. Under these conditions, TfcP is important for PilY1.1 stability. However, at concentrations ≥1 mM added CaCl_2_, calcium alone is sufficient to stabilise PilY1.1 independently of TfcP. Second, in the complete absence of calcium, i.e. after addition of the calcium specific chelator BAPTA, TfcP does not stabilise PilY1.1 while TfcP still accumulates. Third, the PilY1.1^D1173A^ variant, which is predicted to bind calcium with reduced affinity, depends on TfcP for stability at 1.0 mM added CaCl_2_, and even at 10 mM CaCl_2_, this protein is non-functional in the absence of TfcP. Thus, TfcP and calcium both function to stabilise PilY1.1. However, while high calcium concentrations alone can stabilise PilY1.1, TfcP cannot stabilise PilY1.1 in the absence of calcium. Altogether, these findings support a model whereby calcium binding by PilY1.1 is the primary determinant for its stability, and in which TfcP stabilises PilY1.1 at low calcium concentrations by chaperoning calcium binding by PilY1.1. The functional outcome of this TfcP-dependent stimulation of calcium binding by PilY1.1 is that PilY1.1 accumulates at low calcium concentrations and is able to support cluster_1-dependent T4aP formation and T4aPdM. Many myxobacteria including *M. xanthus* are found in terrestrial habitats in which calcium concentrations are described to vary from 0.1-1.0 mM at root-soil interfaces, 3.4-14 mM in some soils and as low as 10-150 µM in other soils^52^. We suggest that TfcP is key to enabling PilY1.1-dependent T4aP formation and T4aPdM in the lower range of calcium concentrations. Interestingly, *M. stipitatus* and *Corallococcus coralloides* only have one gene cluster for minor pilins and PilY1, and this cluster encodes a TfcP ortholog (Fig. 1c; Fig. S1a) emphasising the importance of TfcP in T4aPdM in myxobacteria.

TfcP is a periplasmic protein and contains a non-canonical cytochrome *c* domain in which the second axial heme ligand is a Cys residue rather than the more common His and Met residues in canonical cytochromes *c*. Accordingly, TfcP has a very low redox potential of −304 to −320 mV based on two methods. Moreover, TfcP variants unable to bind heme or with altered heme-binding properties are unstable *in vivo*. *M. xanthus* is strictly aerobic and the genome encodes complex I-IV of the electron transport chain^53^. Thus, the low redox-potential of TfcP supports that it is not part of the respiratory chain, which starts with a potential of −320 mV for the redox pair NAD/NADH^54^. Some cytochromes *c* are involved in electron transport across the OM to external electron acceptors; however, these proteins are canonical cytochromes *c*^55^ suggesting that TfcP also does not engage in this type of electron transport. Some *c*-type cytochromes with His/Cys ligation, e.g. the triheme DsrJ of *Allochromatium vinosum*, are involved in dissimilatory sulfur metabolism in which sulfate is used as terminal electron acceptor^56^. Other *c*-type cytochromes with His/Cys ligation have been suggested to have a role in signalling^35, 57^. Because *M. xanthus* does not respire on sulfate, it is unlikely that TfcP would be involved in dissimilatory sulfur metabolism. While we cannot rule out a function of TfcP in signalling, our data support a scenario in which TfcP is a repurposed cytochrome *c* that is no longer involved in electron transport, and in which the covalently bound heme serves a structural function to stabilise TfcP. This “inert” cytochrome *c* then chaperones calcium binding by PilY1.1 at low calcium concentrations. TfcP also differs from canonical cytochromes *c* by having a C-terminal extension. This extension is important for TfcP stability; however, its precise function remains to be uncovered.

PilY1 proteins of *P. aeruginosa*, *Kingella kingae* and *Neisseria gonorrhoeae* bind calcium using an EF-hand-like calcium binding motif in their C-terminal PilY1 domain (Fig. 6b). An Asp to Ala substitution of the C-terminal Asp residue in this motif abolishes calcium binding and renders the proteins non-functional while overall still folding correctly^27, 46, 47^. The corresponding substitution in the D^1165^xDxDNxxD^1173^ motif in PilY1.1 increased its dependency on TfcP and calcium for stability supporting that PilY1.1 binds calcium as described for other PilY1 proteins. However, while the PilY1 variants of *P. aeruginosa*, *K. kingae* and *N. gonorrhoeae* deficient in calcium binding accumulate, PilY1.1 depends on TfcP for stability at low calcium levels. Also, PilY1.1^D1173A^ only accumulates at very low levels even in the presence of calcium and TfcP, suggesting that PilY1.1 binds calcium with a lower affinity than the three other PilY1 proteins. The observation that PilY1.1 is unstable in the absence of TfcP at low calcium concentrations suggest that the two proteins interact directly. However, such an interaction remains to be shown and will be addressed in future experiments. PilY1.1 as well as the four minor pilins of cluster_1 were detected in purified pili as previously shown for the minor pilins and PilY1.3 of cluster_3^10^. By contrast, we did not detect TfcP in purified pili. We previously showed that sfGFP-tagged PilY1.3 and the sfGFP-tagged minor pilin PilW3 of cluster_3 localise polarly, are incorporated into the T4aPM but do not support pilus extension likely because the sfGFP-tag jams the machine by precluding passage of PilY1.3-sfGFP and PilW3-sfGFP through the secretin channel in the OM^10^. sfGFP-tagged TfcP was fully active and did not localise polarly. These observations strongly support that TfcP is neither part of the T4aPM nor of the pilus. They also strengthen the hypothesis that the suggested direct interaction between PilY1.1 and TfcP is transient and occurs in the periplasm before PilY1.1 incorporation into the T4aPM (Fig. 8). The observation that added calcium stabilises PilY1.1 in the absence of TfcP supports that TfcP does not act as a metallochaperone to deliver calcium to PilY1.1. Altogether, we suggest that TfcP transiently interacts with PilY1.1, thereby stimulating folding and efficient calcium binding by PilY1.1. Subsequently, PilY1.1 with bound calcium is incorporated into the priming complex of the T4aPM to support T4aP formation (Fig. 8). This mechanism of protein stabilisation is reminiscent to that of periplasmic chaperones, which in an ATP-independent manner transiently interact with their periplasmic clients to enable folding^58^, except that the TfcP/PilY1.1 interaction is suggested to promote calcium binding by PilY1.1, which then stabilises PilY1.1. Altogether, these findings also provide evidence for a novel cytochrome *c* function in protein folding and/or stabilisation.

In the presence of added calcium at 1.0 mM, more T4aP are formed and the ratio between minor pilins and PilY1.1 to PilA is increased. These observations support that calcium not only helps to stabilise PilY1.1, but may also stabilise the pilus including the minor pilin/PilY1.1 tip complex in extracellular space. In this context, it is interesting to note that calcium binding has been reported to stabilise the interactions between major pseudopilin subunits in the pseudopilus of the type II secretion system of *Klebsiella oxytoca*^59^. In future experiments, this effect of calcium will be addressed.

In addition to the conserved proteins of the T4aPM, T4aP extension in several species depends on accessory factors that are much less conserved. For instance, the c-di-GMP binding protein FimX in *P. aeruginosa* and SgmX in *M. xanthus* stimulate T4aP extension^60–62^. TfcP adds to be list of such regulators and also acts at the level of extension; however, in contrast to these cytoplasmic regulators, TfcP acts in the periplasm.

## Methods

### Bacterial strains and growth media

All *M. xanthus* strains are derivatives of DK1622^63^ and listed in Supplementary Table 1. All plasmids are listed in Supplementary Table 2. In-frame deletion mutants were generated using double homologous recombination using a *galK-* containing plasmid^64^. Genes were ectopically expressed from the *pilA-*promoter in plasmids integrated by site-specific recombination at the *attB* site. All plasmids were verified by sequencing. All strains were confirmed by PCR. Oligonucleotides are listed in Supplementary Table 3. *M. xanthus* liquid cultures were grown in 1% CTT broth (1% Bacto Casitone (Gibco), 10 mM Tris-HCl pH8.0, 1 mM KPO_4_ pH7.6, 8 mM MgSO_4_) or on 1% CTT 1.5% agar plates. When required media were supplemented with kanamycin (50 µg ml^-1^) or oxytetracyclin (10 µg ml^-1^)^65^. *E. coli* strains were grown in LB broth^66^. Plasmids were propagated using *E. coli* NEB-Turbo.

### Bioinformatics

Homologs of TfcP were searched using BlastP^67^. Pairwise sequence alignments were calculated using EMBOSS-Needle^68^. Protein domains were identified using InterPro^69^. Alignments of TfcP and homologs were computed using MUSCLE^68^. The homology model of PilY1.1 was generated using the Phyre2 server^70^.

### Motility assay

T4aPdM was assayed as described^71^. Briefly, exponentially growing *M. xanthus* cultures were harvested and concentrated in 1% CTT to a density of 7×10^9^ cells ml^-1^. 5 µl of the concentrated cell suspension were spotted on soft-agar CTT plates (0.5% CTT, 10 mM Tris-HCl pH 8.0, 1 mM KPO_4_ pH 7.6, 8 mM MgSO_4_, 0.5% select-agar (Invitrogen)) and incubated at 32°C for 24 hrs. Colonies were imaged using a Leica MZ75 stereomicroscope equipped with a Leica MC120 HD camera.

### T4aP shearing assay

T4aP were sheared off *M. xanthus* cells as described^72^. Briefly, cells were grown on CTT 1.5% agar plates at 32°C for three days, then scraped off, and resuspended in pili resuspension buffer (100 mM Tris-HCl pH 7.6, 150 mM NaCl) (1 ml per 60 mg cells). Cell suspensions were vortexed 10 min at maximum speed. A 100 µl aliquot was harvested and resuspended in 200 µl sodium dodecyl sulfate (SDS) lysis buffer (50 mM Tris-HCl pH 6.8, 2% SDS, 10% glycerol, 0.1 M DTT, 1.5 mM EDTA, 0.001% Bromophenol Blue), and denatured at 95°C for 10 min and used to determine the cellular PilA amount. The remaining cell suspension was cleared three times by 20 min centrifugation at 20,000 *g* at 4°C. Pili in the cleared supernatant were precipitated by adding 10× pili-precipitation buffer (final concentration: 100 mM MgCl_2_, 2% PEG 6000, 100 mM Tris-HCl pH 7.6, 150 mM NaCl), incubation on ice for 4 hrs and centrifugation at 20,000 *g* for 30 min, 4°C. The pellet was resuspended in 1 µl SDS lysis buffer per mg cells and boiled for 10 min at 95°C. The samples were separated by SDS-PAGE and analysed for PilA accumulation by immuno-blot using PilA antibodies.

### Immuno-blot and peroxidase staining

Immuno-blot analysis was carried out as described^66^. Samples were prepared by harvesting exponentially growing *M. xanthus* cells and subsequently resuspension in SDS lysis buffer to an equal concentration of cells. Immuno-blot was done using as primary antibodies α-PilB, α-PilC, α-PilQ^36^, α-PilA, α-LonD^10^, α-FLAG (Rockland; 600-401-383), α-GFP (Roche; 11814460001), α-MalE (New England Biolabs), α-PilY1.1^73^ and α-TfcP. Antibodies against TfcP were generated by Eurogentec against TfcP^Δ1-18^-His_6_ purified from *E. coli* Rosetta 2(DE3) containing plasmid pMH6 using native Ni-NTA affinity purification. As secondary antibodies, goat α-rabbit immunoglobulin G peroxidase conjugate (Sigma-Aldrich, A8275) and sheep α-mouse immunoglobulin G peroxidase conjugate (Amersham, NXA931) were used. Antibodies and conjugates were used in the following dilutions: 1:500 α-TfcP; 1:1000 α-PilY1.1; 1:3000 α-PilB; α-PilC; 1:5000 α-PilQ; α-PilA; 1:6000 α-LonD; 1:2000 α-GFP, α-MalE, α-FLAG and α-mouse peroxidase conjugate; and, 1:10,000 α-rabbit peroxidase conjugate. Blots were developed using Luminata`™`Western HRP substrate (Millipore). Unless otherwise noted, protein from 3×10^8^ cells were loaded per lane. For peroxidase staining, protein was separated by SDS-PAGE, blotted on a nitrocellulose membrane and developed using Luminata`™`Western HRP substrate.

### Fractionation of *M. xanthus*

*M. xanthus* was fractionated into fractions enriched for soluble, IM and OM proteins as described^74^. Briefly, an exponentially growing *M. xanthus* culture was harvested and the pellet resuspended in lysis buffer (50 mM Tris-HCl pH7.6, Protease inhibitor cocktail (Roche)) (1 ml per 80 mg cells). A 75 µl aliquot was taken as the whole cell sample, suspended with SDS-lysis buffer and boiled 10 min at 95°C. Cells were lysed using sonication and lysates cleared by centrifugation at 8000 *g* for 1 min. The cleared lysate was subjected to ultra-centrifugation using an Air-Fuge (Beckman) at ∼150,000 *g* for 1 hr. The resulting supernatant contains soluble proteins and was mixed with SDS-lysis buffer. The pellet was resuspended in detergent-lysis buffer (50 mM Tris-HCl pH 7.6, 2% Triton X-100) and subjected to ultra-centrifugation as described. The resulting supernatant is enriched for IM proteins while the pellet is enriched for OM proteins. The supernatant was mixed with SDS lysis buffer and the pellet resuspended in SDS-lysis buffer. The samples were analysed by SDS-PAGE and immuno-blot.

### Fluorescence microscopy

Exponentally growing *M. xanthus* cells were spotted on 1% agarose pads supplemented with TPM (10 mM Tris-HCl pH 8.0, 1 mM KPO_4_ pH 7.6, 8 mM MgSO_4_) and incubated for 30 min at 32°C before microscopy. Cells were imaged using a Leica DMI600B microscope with a Hamamatsu Flash 4.0 camera. Images were recorded with Leica MM AF software and processed with Metamorph.

### Targeted proteomics

To identify peptides of T4aPM proteins suitable for targeted-mass spectrometry (MS) analysis, we performed sample preparation on *M. xanthus* cell pellets for total proteome analysis as described^10^. Briefly, proteins were extracted from cell pellets by heat exposure in the presence of 2% sodium-lauroylsarcosinate. Extracts were then reduced, alkylated and digested overnight using trypsin (Promega). Peptides were purified using C18 solid phase extraction and analysed on a Q-Exactive Plus mass spectrometer connected to an Ultimate 3000 RSLC and a nanospray flex ion source (all Thermo Scientific). The peptides were analysed using data dependent acquisition with settings as described^10^. MS raw data were searched using Mascot (Matrix Science) and loaded into Scaffold 4 (Proteome software) for further data evaluation. Peptides considered most amenable for targeted MS were chosen for reference peptide synthesis (JPT Peptide Technologies, Berlin) containing heavy labelled (^13^C and ^15^N) C-terminal Lys or Arg residues with a resulting mass shift of +8 Da and +10 Da, respectively. Sequences of reference peptides are listed in Supplementary Table 4. For targeted MS experiments, reference peptides and iRT retention calibration peptides (Biognosys) were spiked into the *M. xanthus* total proteome peptide samples (generated as described), and analysed by liquid chromatography (LC)-MS.

Peptides were separated on a 90 min gradient from 2-50% acetonitrile at a flow rate of 300 nl min^-1^, and analysed by MS in targeted parallel reaction monitoring (PRM) mode. The mass spectrometer first acquired a full MS-Selected Ion Monitoring (SIM) scan with an MS1 resolution of 70,000, AGC (automatic gain control) target setting of 1e^6^ and 100 ms max injection time. Then PRM scans were carried out with a MS2 resolution of 35,000, AGC target setting of 2e^5^, 100 ms maximum injection time with a quadrupole isolation window of 1.6 m/z. Normalised collision energy was set to 27%. All stages of targeted MS data analysis was carried out in Skyline (20.2.1.384)^75^. Results with dot-product <0.85 or ratio_heavy/light_<0.005 were excluded from the analysis.

### Proteome analysis of T4aP

Label-free quantification (LFQ) MS of the pili proteome was carried out as described^10^. Briefly, pili were purified following the shearing assay protocol with the modification that after precipitation, pili were resuspended in pili-resuspension buffer and re-precipitated with pili-precipitation buffer three times. Pili were resuspended in pili-resuspension buffer to 1 µl buffer per 1 mg cells. 25% of the pili sample was mixed with SDS-lysis buffer and analysed by SDS-PAGE and subsequent staining with Coomassie Blue. The remaining 75% were precipitated with acetone. The dried acetone pellets were resuspended, reduced, alkylated and digested with trypsin as described^10^. Pili LFQ proteomics analysis was carried out on an Exploris 480 mass spectrometer (Thermo Scientific), connected to an Ultimate 3000 RSLC. Peptides were separated on a 60 min gradient from 2-50% acetonitrile at a flow rate of 300 nl min^-1^. The Exploris 480 mass spectrometer first acquired a full MS scan with an MS1 resolution of 60,000, AGC target setting of 3e^6^ and 60 ms max injection time, followed by MS/MS scans of Top-20 most abundant signals. For MS/MS scans a resolution of 7,500 was set, with an AGC of 2e^5^ and 30 ms max. injection time. Normalised collision energy was set to 27% and the isolation window of the quadrupole was 1.6 m/z. All MS raw data was analysed by MaxQuant (1.6.17.0). iBAQ values were calculated as described^10^ as the sum of all peptide intensities for a given protein divided by the number of theoretically MS observable peptides. Following MaxQuant analysis, the iBAQ values were normalised by the total iBAQ sum independently of the highly abundant PilA.

### Purification of MalE-TfcP

For purification of MalE-TfcP/MalE-TfcP^C91M^, gene expression was done in *E. coli* strain BL21 containing the helper plasmid pEC86, which encodes the *ccm* genes for cytochrome *c* maturation of *E. coli*, as well as pMH31 (MalE-TfcP) or pMH39 (MalE-TfcP^C91M^) using auto-induction in buffered 5052-Terrific-Broth (0.5% glycerol, 0.05% glucose, 0.2% lactose, 2.4% yeast extract, 2% tryptone, 25 mM Na_2_HPO_4_, 25 mM KH_2_PO_4_, 50 mM NH_4_Cl, 5 mM Na_2_SO_4_, 2 mM MgSO_4_)^76^ containing chloramphenicol (25 µg ml^-1^) and carbenicillin (100 µg ml^-1^). After 24 hrs incubation at 37°C, cells were harvested, and resuspended in MBP-lysis buffer (100 mM Tris-HCl pH 7.0, 200 mM NaCl) supplemented with EDTA-free protease inhibitor cocktail (Roche) and lysed using sonication. The lysate was cleared by centrifugation at 20,000 *g*, 4°C for 30 minutes and loaded onto a 5 ml HighTrap MBP column (GE Healtcare) using an Äkta-Pure system (GE Healthcare). The column was washed with lysis buffer and protein eluted with 10 column volumes MBP-elution buffer (100 mM Tris-HCl pH 7.0, 200 mM NaCl, 10 mM maltose). The elution fractions containing MalE-TfcP/MalE-TfcP^C91M^ were pooled and diluted four fold in 100 mM Tris-HCl pH 7.0. The pooled and diluted samples were loaded onto a HighTrap SP ion exchange column. The column was washed with IEX-wash buffer (100 mM Tris-HCl pH 7.0) and protein eluted in a linear gradient with IEX-elution buffer (100 mM Tris-HCl pH 7.0, 2 M NaCl). Samples were concentrated using an Amicon Ultra filter with 10 kDa cutoff and loaded on a HiLoad 16/600 Superdex 200 pg (GE Healthcare) size exclusion chromatography column equilibrated with SEC-buffer (50 mM Tris-HCl pH 7.6, 50 mM NaCl). Protein was either used fresh or snap-frozen in buffer containing SEC-buffer with 10% glycerol.

### UV-Vis spectroscopy

UV-Vis measurements of purified (oxidised) and reduced MalE-TfcP/MalE-TfcP^C91M^ was conducted on a Tecan M200Pro platereader or a Shimadzu 1900 spectrophotometer. Protein was diluted to an absorbance of ∼0.7. After measurement of the oxidised spectrum, protein was reduced by adding a few crystals of sodium-dithionite, equilibrated for 15 min and the reduced spectrum recorded.

### Redox titrations

Redox titrations were carried out in a Coy anaerobic tent (3% H_2_, <5 ppm O_2_). MalE-TfcP in HEPES buffer, pH 7.0, was mixed with 20 µM (final concentration) of the following redox mediators: Phenosafranin, safranin T, neutral red, benzyl viologen, and methyl viologen. The solution potential was measured with an InLab redox micro combination electrode (Mettler Toledo) under anaerobic conditions. Correction to redox potentials vs. H_2_/H^+^ was done by addition of 207 mV to the reading of the potentiometer. Stirring was done using a 8 mm teflon coated stirrer bar. For redox titration using visible spectroscopy (using a Shimadzu 1900 spectrophotometer), automated addition of 15 µl buffered 0.2 mM sodium-dithionite solution was done using a remotely controlled peristaltic pump (Pharmacia P1) for 60 sec followed by 2 min equilibration and 2 min recording of the spectra in the 600-460 nm range. The normalised absorbance increase at 550 nm (corrected by the absorbance for titration of mediators only) was fitted to the Nernst equation for n=1 at 298 K. For the EPR titration, manual addition of aliquots of buffered sodium-dithionite was used. After stabilisation of the solution potential, 300 µl samples were withdrawn, transferred to EPR tubes, which were capped with rubber tubing and an acrylic glass stick. Samples were shock-frozen and stored in liquid nitrogen until the EPR measurements.

### EPR spectroscopy

EPR spectra were recorded with an X-band EPR spectrometer (Bruker Elexsys E580) in a 4122HQE-W1/1017 resonator. The temperature of the samples in Ilmasil PN quartz tubes (4.7±0.2 mm outer diameter, 0.45±0.05 mm wall thickness) was maintained at 12 K with an ESR900 continuous flow helium cryostat (Oxford Instruments). The modulation frequency was 100 kHz and the modulation amplitude 1.5 mT. Spectra were averages for four 90 sec scans. For the titration, the normalised amplitude of the derivative-shaped feature of the low spin EPR signal of the ferric state at *g*=2.26 was used for a fit to the Nernst equation (n=1, T=298 K).

### Operon mapping

Total RNA was isolated from exponentially growing *M. xanthus* cultures using the Monarch Total RNA Miniprep Kit (NEB). 10^9^ cells were harvested and resuspended in 200 µl lysis-buffer (100 mM Tris-HCl pH 7.6, 1 mg ml^-1^ lysozyme). After incubation at 25°C for 5 min cells were lysed and RNA purified according to manufacturer’s protocol with the exception that the on-column DNase treatment was omitted. RNA was eluted in RNase-free water and subsequently treated with Turbo DNase and purified using the Monarch RNA Cleanup Kit (50 µg) (NEB) and eluted in RNase-free water. 1 µg of RNA was used for cDNA synthesis using the LunaScript RT SuperMix Kit (NEB) with and without reverse transcriptase (RT). cDNA was diluted 1:5 with water and 1 µl of diluted cDNA used for PCR reactions.

### qRT-PCR

For qRT-PCR RNA was isolated and cDNA synthesised as described for operon mapping. qPCRs were carried out using the Luna Universal qPCR MasterMix (NEB) with the primers listed in Supplementary Table 3 and measured on an Applied Biosystems 7500 Real-Time PCR system. Relative gene expression levels were calculated using the comparative C_T_ method^77^. *Mxan_3298* (*tuf2*), which encodes elongation factor Tu, and *mxan_3303* (*rpsS*), which encodes the small ribosomal subunit protein S19, were used as internal controls. All experiments were done with three biological replicates and two technical replicates.

### Statistics and reproducibility

Data shown for operon mapping, T4aP-dependent motility, T4aP shearing assays, immuno-blot experiments, UV-Vis spectroscopy and fluorescence microscopy were obtained in at least two biological replicates with similar results. For targeted proteomics and LFQ-analysis of the pili proteome, four biological replicates were analysed. qRT-PCR analysis were conducted with three biological replicates each with two technical replicates. Redox titrations and EPR-spectroscopy where done in a single experiment.

### Data availability

Source data are provided with this paper. The authors declare that all data supporting this study are available within the article, its Supplementary Information file, and the Source Data file.

## Supporting information

All supplemental information

## Acknowledgements

We thank Steffi Lindow for excellent help with plasmid and strain constructions, Bazlur Rashid for help with preparation of the EPR samples, and Seigo Shima as well as Rolf Thauer for many helpful discussions.

## Authors’ contributions

MH: Designed and conceived the study and performed most of the experiments. ATL: Conceived the study, supervised research and provided strains and plasmids. TG: Performed targeted and label-free mass spectrometry-based quantitative proteomics. NW: Helped with the *in vitro* analyses of MalE-TfcP. EH: Generated the PilY1.1 antibodies. AJP: Conceived and supervised the *in vitro* analyses of MalE-TfcP. LSA: Conceived the study, supervised research and provided funding. MH, ATL, TG, AJP and LSA: Analysed and interpreted data and wrote the manuscript.

All authors approved the final manuscript.

## References

1 Harshey, R. M. Bacterial motility on a surface: many ways to a common goal. Annu. Rev Microbiol. 57, 249–273 (2003).

2 Burrows, L. L. *Pseudomonas aeruginosa* twitching motility: type IV pili in action. Annu. Rev. Microbiol. 66, 493–520 (2012).

3 Evans, K. J., Lambert, C. & Sockett, R. E. Predation by *Bdellovibrio bacteriovorus* HD100 requires type IV pili. J. Bacteriol. 189, 4850–4859 (2007).

4 Craig, L., Forest, K. T. & Maier, B. Type IV pili: dynamics, biophysics and functional consequences. Nat. Rev. Microbiol. 17, 429–440 (2019).

5 Merz, A. J., So, M. & Sheetz, M. P. Pilus retraction powers bacterial twitching motility. Nature 407, 98–102 (2000).

6 Skerker, J. M. & Berg, H. C. Direct observation of extension and retraction of type IV pili. Proc Natl Acad Sci U S A 98, 6901–6904 (2001).

7 Clausen, M., Jakovljevic, V., Søgaard-Andersen, L. & Maier, B. High force generation is a conserved property of type IV pilus systems. J. Bacteriol. 191, 4633–4638 (2009).

8 Chang, Y. W. et al. Architecture of the type IVa pilus machine. Science 351, aad2001 (2016).

9 Gold, V. A., Salzer, R., Averhoff, B. & Kühlbrandt, W. Structure of a type IV pilus machinery in the open and closed state. eLife 4, e07380 (2015).

10 Treuner-Lange, A. et al. PilY1 and minor pilins form a complex priming the type IVa pilus in *Myxococcus xanthus*. Nat. Comm. 11, 5054 (2020).

11 McCallum, M., Tammam, S., Khan, A., Burrows, L. L. & Howell, P. L. The molecular mechanism of the type IVa pilus motors. Nat. Comm. 8, 15091 (2017).

12 Jakovljevic, V., Leonardy, S., Hoppert, M. & Søgaard-Andersen, L. PilB and PilT are ATPases acting antagonistically in type IV pilus function in *Myxococcus xanthus*. J. Bacteriol. 190, 2411–2421 (2008).

13 Mancl, J. M., Black, W. P., Robinson, H., Yang, Z. & Schubot, F. D. Crystal structure of a type IV pilus assembly ATPase: Insights into the molecular mechanism of PilB from *Thermus thermophilus*. Structure 24, 1886–1897 (2016).

14 Nguyen, Y. et al. *Pseudomonas aeruginosa* minor pilins prime type IVa pilus assembly and promote surface display of the PilY1 adhesin. J. Biol. Chem. 290, 601–611 (2015).

15 Misic, A. M., Satyshur, K. A. & Forest, K. T. *P. aeruginosa* PilT structures with and without nucleotide reveal a dynamic type IV pilus retraction motor. J. Mol. Biol. 400, 1011–1021 (2010).

16 Rudel, T., Scheurerpflug, I. & Meyer, T. F. *Neisseria* PilC protein identified as type-4 pilus tip-located adhesin. Nature 373, 357–359 (1995).

17 Johnson, M. D. et al. *Pseudomonas aeruginosa* PilY1 binds integrin in an RGD-and calcium-dependent manner. PLOS One 6, e29629 (2011).

18 Marko, V. A., Kilmury, S. L. N., MacNeil, L. T. & Burrows, L. L. *Pseudomonas aeruginosa* type IV minor pilins and PilY1 regulate virulence by modulating FimS-AlgR activity. PLoS Pathogens 14, e1007074 (2018).

19 Kuchma, S. L. et al. Cyclic-di-GMP-mediated repression of swarming motility by *Pseudomonas aeruginosa*: the *pilY1* gene and its impact on surface-associated behaviors. J. Bacteriol. 192, 2950–2964 (2010).

20 Pelicic, V. Type IV pili: e pluribus unum? Mol. Microbiol. 68, 827–837 (2008).

21 Zollner, R., Cronenberg, T. & Maier, B. Motor properties of PilT-independent type 4 pilus retraction in *Gonococci*. J. Bacteriol. 201 (2019).

22 Feng, T. et al. Interspecies and intraspecies signals synergistically regulate *Lysobacter enzymogenes* twitching motility. Appl. Env. Microbiol. 85 (2019).

23 Cruz, L. F., Parker, J. K., Cobine, P. A. & De La Fuente, L. Calcium-enhanced twitching motility in *Xylella fastidiosa* is linked to a single PilY1 homolog. App. Env. Microbiol. 80, 7176–7185 (2014).

24 Engel, C. E. A., Vorlander, D., Biedendieck, R., Krull, R. & Dohnt, K. Quantification of microaerobic growth of *Geobacter sulfurreducens*. PLoS One 15, e0215341 (2020).

25 Parker, J. K., Cruz, L. F., Evans, M. R. & De La Fuente, L. Presence of calcium-binding motifs in PilY1 homologs correlates with Ca-mediated twitching motility and evolutionary history across diverse bacteria. FEMS Microbiol. Lett. 362 (2015).

26 Giltner, C. L., Nguyen, Y. & Burrows, L. L. Type IV pilin proteins: versatile molecular modules. Microbiol. Mol. Biol. Rev. 76, 740–772 (2012).

27 Orans, J. et al. Crystal structure analysis reveals *Pseudomonas* PilY1 as an essential calcium-dependent regulator of bacterial surface motility. Proc. Natl. Acad Sci. U S A 107, 1065–1070 (2010).

28 Hoppe, J. et al. PilY1 promotes *Legionella pneumophila* infection of human lung tissue explants and contributes to bacterial adhesion, host cell invasion, and twitching motility. Front. Cell. Infect. Microbiol. 7, 63 (2017).

29 Zhang, Y., Ducret, A., Shaevitz, J. & Mignot, T. From individual cell motility to collective behaviors: insights from a prokaryote, *Myxococcus xanthus*. FEMS Microbiol. Rev. 36, 149–164 (2012).

30 Schumacher, D. & Søgaard-Andersen, L. Regulation of cell polarity in motility and cell division in *Myxococcus xanthus*. Annu. Rev. Microbiol. 71, 61–78 (2017).

31 Zaidi, S., Hassan, M. I., Islam, A. & Ahmad, F. The role of key residues in structure, function, and stability of cytochrome-*c*. Cell. Mol. Life Sci. 71, 229–255 (2014).

32 Thöny-Meyer, L. Cytochrome c maturation: a complex pathway for a simple task? Biochem. Soc. Trans. 30, 633–638 (2002).

33 Ambler, R. P. Sequence variability in bacterial cytochromes c. Biochim Biophys Acta 1058, 42–47 (1991).

34 Fufezan, C., Zhang, J. & Gunner, M. R. Ligand preference and orientation in *b*-and *c*-type heme-binding proteins. Proteins: Structure, Function, and Bioinformatics 73, 690–704 (2008).

35 Zuccarello, L. et al. Protonation of the Cysteine axial ligand investigated in His/Cys *c*-type cytochrome by UV-Vis and Mid-and Far-IR spectroscopy. J. Phys. Chem. Lett. 11, 4198–4205 (2020).

36 Bulyha, I. et al. Regulation of the type IV pili molecular machine by dynamic localization of two motor proteins. Mol. Microbiol. 74, 691–706 (2009).

37 Friedrich, C., Bulyha, I. & Søgaard-Andersen, L. Outside-in assembly pathway of the type IV pilus system in *Myxococcus xanthus*. J. Bacteriol. 196, 378–390 (2014).

38 Nudleman, E., Wall, D. & Kaiser, D. Polar assembly of the type IV pilus secretin in *Myxococcus xanthus*. Mol. Microbiol. 60, 16–29 (2006).

39 Londer, Y. Y. in Heterologous gene expression in E. coli: Methods and protocols Vol. 705 123–150 (Humana Press, 2011).

40 Poulos, T. L. Heme enzyme structure and function. Chem. Rev. 114, 3919–3962 (2014).

41 Raphael, A. L. & Gray, H. B. Axial ligand replacement in horse heart cytochrome *c* by semisynthesis. Proteins 6, 338–340 (1989).

42 Raphael, A. L. & Gray, H. B. Semisynthesis of axial-ligand (position 80) mutants of cytochrome *c*. J. Am. Chem. Soc. 113, 1038–1040 (2002).

43 Smith, A. T. et al. Identification of Cys 94 as the distal ligand to the Fe (III) heme in the transcriptional regulator RcoM-2 from *Burkholderia xenovorans*. J. Biol. Inorg. Chem. 17, 1071–1082 (2012).

44 Cheesman, M. R., Little, P. J. & Berks, B. C. Novel heme ligation in a *c*-type cytochrome involved in thiosulfate oxidation: EPR and MCD of SoxAX from *Rhodovulum sulfidophilum*. Biochemistry 40, 10562–10569 (2001).

45 Reijerse, E. J. et al. The unusal redox centers of SoxXA, a novel c-type heme-enzyme essential for chemotrophic sulfur-oxidation of *Paracoccus pantotrophus*. Biochemistry 46, 7804–7810 (2007).

46 Cheng, Y. et al. Mutation of the conserved calcium-binding motif in *Neisseria gonorrhoeae* PilC1 impacts adhesion but not piliation. Infect. Imm. 81, 4280–4289 (2013).

47 Porsch, E. A. et al. Calcium binding properties of the *Kingella kingae* PilC1 and PilC2 proteins have differential effects on type IV pilus-mediated adherence and twitching motility. J. Bacteriol. 195, 886–895 (2013).

48 Kuner, J. M. & Kaiser, D. Fruiting body morphogenesis in submerged cultures of *Myxococcus xanthus*. J Bacteriol 151, 458–461 (1982).

49 Sun, M., Wartel, M., Cascales, E., Shaevitz, J. W. & Mignot, T. Motor-driven intracellular transport powers bacterial gliding motility. Proc. Natl. Acad Sci. U S A 108, 7559–7564 (2011).

50 Nan, B. et al. Myxobacteria gliding motility requires cytoskeleton rotation powered by proton motive force. Proc. Natl. Acad. Sci. U S A 108, 2498–2503 (2011).

51 Bilecen, K. & Yildiz, F. H. Identification of a calcium-controlled negative regulatory system affecting *Vibrio cholerae* biofilm formation. Env. Microbiol. 11, 2015–2029 (2009).

52 McLaughlin, S. B. & Wimmer, R. Tansley Review No. 104. Calcium physiology and terrestrial ecosystem processes. New Phytol. 142, 373–417 (1999).

53 Goldman, B., Bhat, S. & Shimkets, L. J. Genome evolution and the emergence of fruiting body development in *Myxococcus xanthus*. PLoS One 2, e1329 (2007).

54 Kracke, F., Vassilev, I. & Krömer, J. O. Microbial electron transport and energy conservation–the foundation for optimizing bioelectrochemical systems. Front. Microbiol. 6, 575 (2015).

55 Edwards, M. J., White, G. F., Butt, J. N., Richardson, D. J. & Clarke, T. A. The crystal structure of a biological insulated transmembrane molecular wire. Cell 181, 665–673 e610 (2020).

56 Grein, F. et al. DsrJ, an essential part of the DsrMKJOP transmembrane complex in the purple sulfur bacterium *Allochromatium vinosum*, is an unusual triheme cytochrome *c*. Biochemistry 49, 8290–8299 (2010).

57 Shimizu, T. Binding of cysteine thiolate to the Fe(III) heme complex is critical for the function of heme sensor proteins. J. Inorg. Biochem. 108, 171–177 (2012).

58 Goemans, C., Denoncin, K. & Collet, J. F. Folding mechanisms of periplasmic proteins. Biochim. Biophys. Acta 1843, 1517–1528 (2014).

59 López-Castilla, A. et al. Structure of the calcium-dependent type 2 secretion pseudopilus. Nat. Microbiol. 2, 1686–1695 (2017).

60 Jain, R., Sliusarenko, O. & Kazmierczak, B. I. Interaction of the cyclic-di-GMP binding protein FimX and the Type 4 pilus assembly ATPase promotes pilus assembly. PLOS Pathogens 13, e1006594 (2017).

61 Potapova, A., Carreira, L. A. M. & Søgaard-Andersen, L. The small GTPase MglA together with the TPR domain protein SgmX stimulates type IV pili formation in *M. xanthus*. Proc. Natl. Acad Sci. U S A 117, 23859–23868 (2020).

62 Mercier, R. et al. The polar Ras-like GTPase MglA activates type IV pilus via SgmX to enable twitching motility in *Myxococcus xanthus*. Proc. Natl. Acad Sci. U S A 117, 28366–28373 (2020).

63 Kaiser, D. Social gliding is correlated with the presence of pili in *Myxococcus xanthus*. Proc. Natl. Acad. Sci. U S A 76, 5952–5956 (1979).

64 Shi, X. et al. Bioinformatics and experimental analysis of proteins of two-component systems in *Myxococcus xanthus*. J. Bacteriol. 190, 613–624 (2008).

65 Søgaard-Andersen, L., Slack, F. J., Kimsey, H. & Kaiser, D. Intercellular C-signaling in *Myxococcus xanthus* involves a branched signal transduction pathway. Genes Dev. 10, 740–754 (1996).

66 Sambrook, J., Fritsch, E. F. & Maniatis, T. Molecular Cloning: A Laboratory Manual. (Cold Spring Harbor Laboratory Press), (1989).

67 Altschul, S. F., Gish, W., Miller, W., Myers, E. W. & Lipman, D. J. Basic local alignment search tool. J. Mol. Biol. 215, 403–410 (1990).

68 Madeira, F., et al. The EMBL-EBI search and sequence analysis tools APIs in 2019. Nucl. Acids Res. 47, W636–W641 (2019).

69 Blum, M. et al. The InterPro protein families and domains database: 20 years on. Nucl. Acids Res. 49, D344–D354 (2021).

70 Kelley, L. A., Mezulis, S., Yates, C. M., Wass, M. N. & Sternberg, M. J. The Phyre2 web portal for protein modeling, prediction and analysis. Nat. Protoc. 10, 845–858 (2015).

71 Shi, W. & Zusman, D. R. The two motility systems of *Myxococcus xanthus* show different selective advantages on various surfaces. Proc Natl Acad Sci U S A 90, 3378–3382 (1993).

72 Wu, S. S., Wu, J. & Kaiser, D. The *Myxococcus xanthus pilT* locus is required for social gliding motility although pili are still produced. Mol. Microbiol. 23, 109–121 (1997).

73 Agrawal, S. Identification and characterization of novel factors needed for two aspects of Myxococcus xanthus physiology: Social motility and osmoregulation. PhD thesis, Johns Hopkins University, (2013).

74 Thomasson, B. et al. MglA, a small GTPase, interacts with a tyrosine kinase to control type IV pili-mediated motility and development of *Myxococcus xanthus*. Mol. Microbiol. 46, 1399–1413 (2002).

75 MacLean, B. et al. Skyline: an open source document editor for creating and analyzing targeted proteomics experiments. Bioinformatics 26, 966–968 (2010).

76 Studier, F. W. Stable expression clones and auto-induction for protein production in *E. coli*. Methods Mol. Biol. 1091, 17–32 (2014).

77 Schmittgen, T. D. & Livak, K. J. Analyzing real-time PCR data by the comparative CT method. Nat. Protoc. 3, 1101–1108 (2008).

