## Supplementary material for "A repurposed, non-canonical cytochrome *c*, chaperones calcium binding by PilY1 for type IVa pili formation": All supplemental information

**This file contains:**

- Supplementary Figures 1-4
- Supplementary Tables 1-4
- Supplementary References

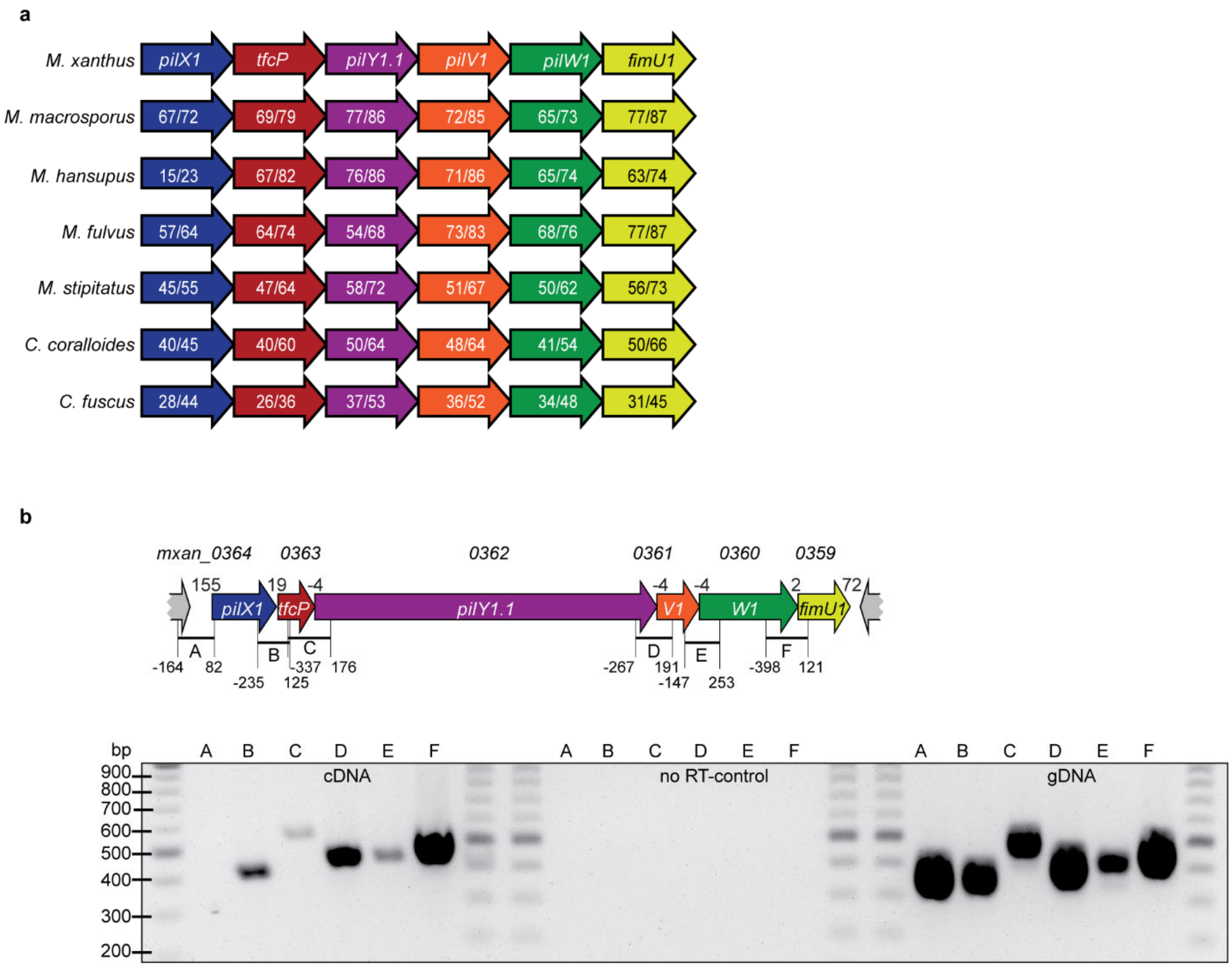


**Figure S1. Cluster_1 is conserved in myxobacteria.**
**a** Comparison of cluster_1 homologs in myxobacteria. All six proteins in the listed species are encoded at the same locus. Arrows indicate direction of transcription. Numbers within genes represent identity/similarity determined by pairwise alignment with the respective *M. xanthus* protein. **b** Operon mapping of cluster_1 in *M. xanthus*. Upper panel, genetic organization of cluster_1. Locus tags are included above genes and gene names within genes. Distances between start and stop codons are shown above. Letters below arrows indicate the fragments amplified by PCR. Numbers indicate the distance from the 5'-end of a primer to the first base of the stop codon or the first base of the start codon as appropriate. The PCR products amplified using cDNA, an enzyme free reverse transcription reaction and genomic DNA as templates were separated on a 1% agarose gel. Letters above the individual lanes correspond to the letters of the primer combinations depicted above. Molecular size markers in base-pairs are shown on the left.

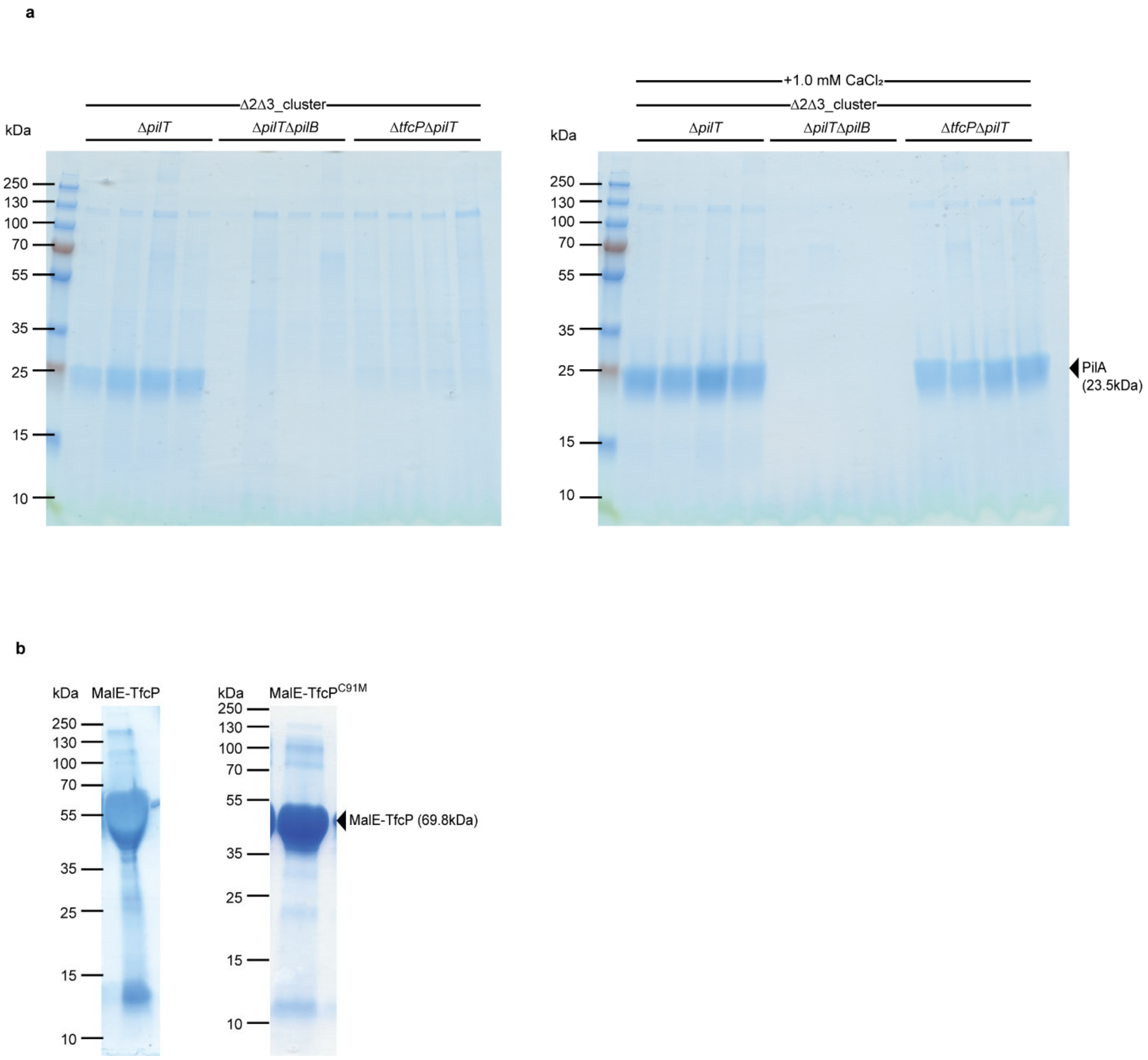


**Figure S2. Purified pili and MalE-TfcP variants**

**a** Purified pili used for proteome analysis. Pili from 15 mg of cells were loaded on SDS-PAGE and stained with Coomassie Blue. The left and right gels show pili from cells grown on 1.5% agar supplemented with 1.0%CTT in the absence and presence of 1.0 mM additional $CaCl_2$, respectively. **b** Purified MalE-TfcP variants used for spectroscopic analysis. ~5 µg of purified proteins were separated by SDS-PAGE and stained by Coomassie Blue.

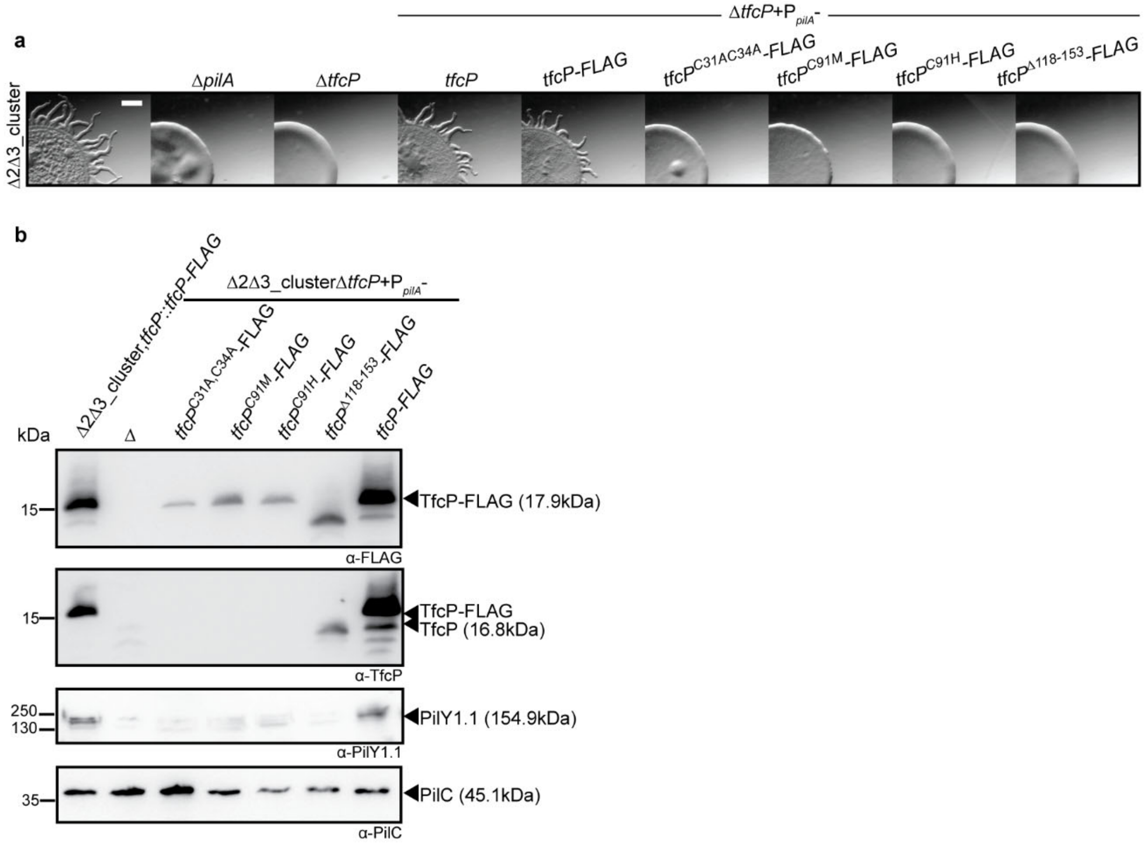


**Figure S3. Amino acid substitution in heme-binding residues of TfcP or deletion of the C-terminal extension affect protein stability**

**a** Assays for T4aPdM. $WT_{\Delta 2\Delta 3}$ and strains expressing mutant TfcP variants were spotted on 0.5% agar supplemented with 0.5% CTT and imaged after 24 hrs. Scale bar, 1 mm. **b** Accumulation of TfcP and PilY1.1 in strains expressing TfcP variants. Protein from the same number of cells grown in 1.0% CTT suspension culture was separated by SDS-PAGE and analysed by immuno-blotting. The lane labeled with Δ contains whole cell lysate of a Δ1Δ2_cluster strain as a negative control. PilC was used as a loading control.

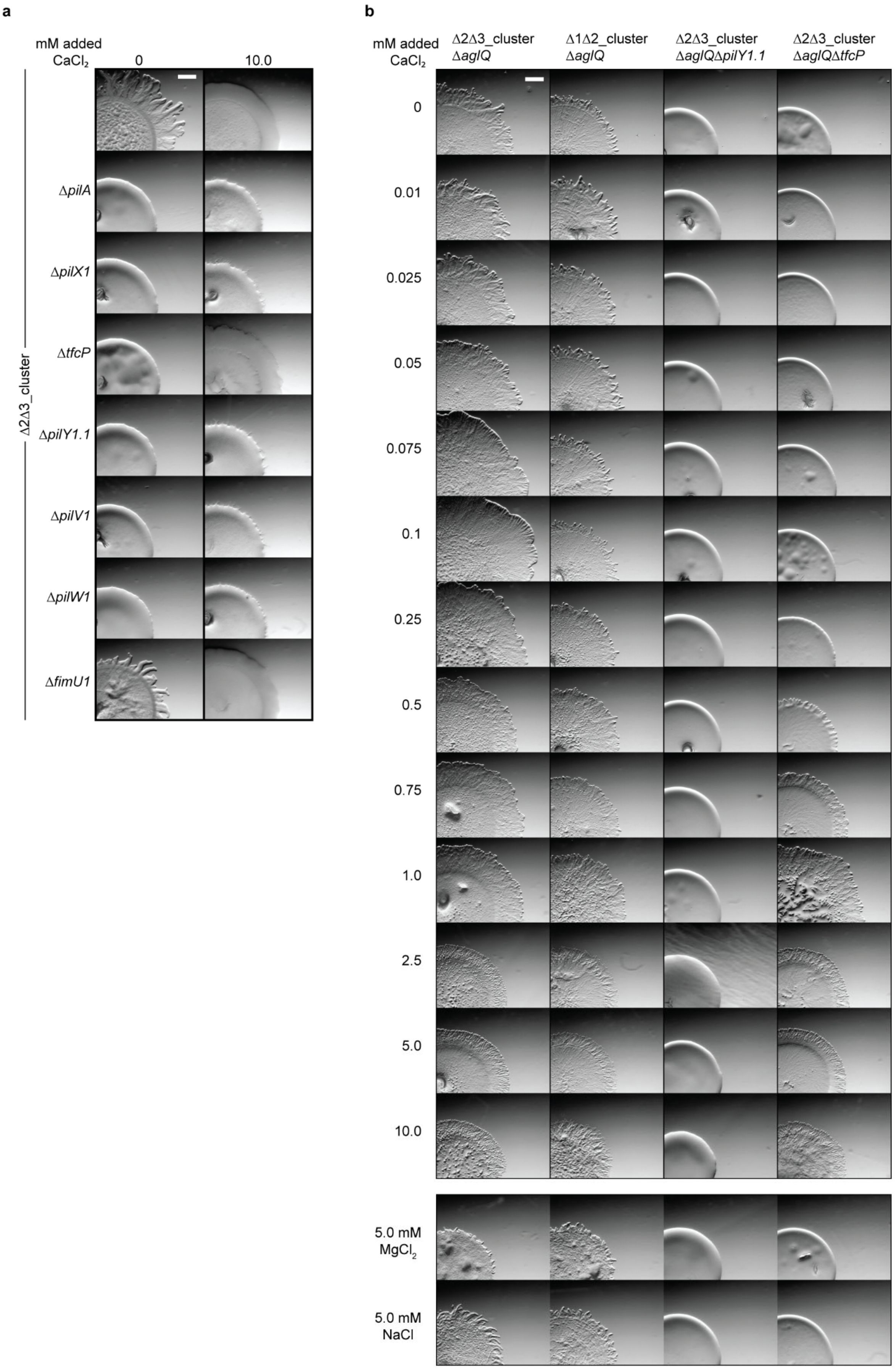
a
mM added CaCl$_2$
0
10.0
Δ2Δ3_cluster
Δ*pilA*
Δ*pilX1*
Δ*tfcP*
Δ*pilY1.1*
Δ*pilV1*
Δ*pilW1*
Δ*fimU1*
b
mM added CaCl$_2$
Δ2Δ3_cluster Δ*aglQ*
Δ1Δ2_cluster Δ*aglQ*
Δ2Δ3_cluster Δ*aglQ*Δ*pilY1.1*
Δ2Δ3_cluster Δ*aglQ*Δ*tfcP*
0
0.01
0.025
0.05
0.075
0.1
0.25
0.5
0.75
1.0
2.5
5.0
10.0
5.0 mM MgCl$_2$
5.0 mM NaCl

**Figure S4. $CaCl_2$ affects T4aPdM of *M. xanthus***

**a** Assay for T4aPdM. Cells were grown in 1.0% CTT suspension culture and plated on 0.5% agar supplemented with 0.5% CTT and imaged after 24 hrs. The final concentration of added $CaCl_2$ is indicated. Scale bar, 1 mm. **b** Assay for T4aPdM. Cells were grown in 1.0% CTT suspension culture and plated on 0.5% agar supplemented with 0.5% CTT and imaged after 24 hrs. The final concentration of added $CaCl_2$, $MgCl_2$ and NaCl is indicated. Scale bar, 1 mm. The experiments at 0, 0.25, 0.5, 0.75 and 1.0 mM added $CaCl_2$ for the three $WT_{\Delta 2 \Delta 3}\Delta$*aglQ* strains are also included in Fig. 6c and are included for comparison.

**Supplementary Table 1. Strains used in this study**

| Name | Description | Reference |
|---|---|---|
| | ***M. xanthus* strains** | |
| SA6892 | Δcluster_2 (mxan_1021-1017) Δcluster_3 (mxan_1369-1365) | [1] |
| SA6888 | Δcluster_1 (mxan_0364-0359) Δcluster_2 | [1] |
| SA8754 | Δcluster_2 Δcluster_3 ΔpilT (mxan_5787) ΔpilB (mxan_5788) | This study |
| SA7717 | Δcluster_2 Δcluster_3 ΔpilT | This study |
| SA7698 | Δcluster_2 Δcluster_3 ΔaglQ (mxan_6861) | This study |
| SA7703 | Δcluster_1 Δcluster_2 ΔaglQ | This study |
| SA7649 | Δcluster_2 Δcluster_3 ΔpilX1 (mxan_0364) | This study |
| SA7648 | Δcluster_2 Δcluster_3 ΔtfcP (mxan_0363) | This study |
| SA7647 | Δcluster_2 Δcluster_3 ΔpilY1.1 (mxan_0362) | This study |
| SA7646 | Δcluster_2 Δcluster_3 ΔpilV1 (mxan_0361) | This study |
| SA7645 | Δcluster_2 Δcluster_3 ΔpilW1 (mxan_0360) | This study |
| SA7644 | Δcluster_2 Δcluster_3 ΔfimU1 (mxan_0359) | This study |
| SA7672 | Δcluster_2 Δcluster_3 ΔpilT ΔpilX1 | This study |
| SA7673 | Δcluster_2 Δcluster_3 ΔpilT ΔtfcP | This study |
| SA7674 | Δcluster_2 Δcluster_3 ΔpilT ΔpilY1.1 | This study |
| SA7675 | Δcluster_2 Δcluster_3 ΔpilT ΔpilV1 | This study |
| SA7676 | Δcluster_2 Δcluster_3 ΔpilT ΔpilW1 | This study |
| SA7677 | Δcluster_2 Δcluster_3 ΔpilT ΔfimU1 | This study |
| SA9004 | Δcluster_2 Δcluster_3 tfcP::tfcP-FLAG | This study |
| SA9009 | Δcluster_2 Δcluster_3 tfcP::tfcP-sfGFP | This study |
| SA9012 | Δcluster_2 Δcluster_3 ΔpilY1.1 ΔaglQ | This study |
| SA9016 | Δcluster_2 Δcluster_3 ΔtfcP ΔaglQ | This study |
| SA9017 | Δcluster_2 Δcluster_3 ΔpilA (mxan_5783) | This study |
| SA9019 | Δcluster_2 Δcluster_3 ΔtfcP pilM (mxan_5776)::mCherry-pilM | This study |
| SA7680 | Δcluster_2 Δcluster_3 ΔtfcP attB::$P_{pilA}$-tfcP | This study |
| SA9040 | Δcluster_2 Δcluster_3 ΔtfcP attB::$P_{pilA}$ -tfcP-FLAG | This study |
| SA9041 | Δcluster_2 Δcluster_3 ΔtfcP attB::$P_{pilA}$ -tfcP$^{C91H}$-FLAG | This study |
| SA9042 | Δcluster_2 Δcluster_3 ΔtfcP attB::$P_{pilA}$ -tfcP$^{C91M}$-FLAG | This study |
| SA9043 | Δcluster_2 Δcluster_3 ΔtfcP attB::$P_{pilA}$ -tfcP$^{\Delta 118-153}$-FLAG | This study |
| SA9031 | Δcluster_2 Δcluster_3 ΔpilY1.1 attB::$P_{pilA}$ -pilY1.1 | This study |
| SA9032 | Δcluster_2 Δcluster_3 ΔfimU1 attB::$P_{pilA}$ -fimU1 | This study |
| SA9033 | Δcluster_2 Δcluster_3 ΔpilV1 attB::$P_{pilA}$ -pilV1 | This study |
| SA9044 | Δcluster_2 Δcluster_3 ΔtfcP attB::$P_{pilA}$ -tfcP$^{C31A,C34A}$-FLAG | This study |
| SA9034 | Δcluster_2 Δcluster_3 ΔpilW1 attB::$P_{pilA}$ -pilW1 | This study |
| SA9055 | Δcluster_2 Δcluster_3 ΔpilX1 attB::$P_{pilA}$ -pilX1 | This study |
| SA9051 | Δcluster_2 Δcluster_3 pilM::mCherry-pilM | This study |
| SA9064 | Δcluster_2 Δcluster_3 pilY1.1::pilY1.1$^{D1173A}$ | This study |
| SA9066 | Δcluster_2 Δcluster_3 ΔtfcP pilY1.1::pilY1.1$^{D1173A}$ | This study |
| SA9068 | Δcluster_2 Δcluster_3 ΔaglQ pilY1.1::pilY1.1$^{D1173A}$ | This study |

| | | |
|---|---|---|
| SA9069 | Δ*cluster_2* Δ*cluster_3* Δ*tfcP* Δ*aglQ* *pilY1.1::pilY1.1*$^{D1173A}$ | This study |
| SA6024 | Δ*pilBTCMNOPQ* | [2] |
| | ***E. coli*** **strains** | |
| NEB-Turbo | F' *proA*$^{+}$*B*$^{+}$ *lacI*$^{q}$ Δ*lacZM15* / *fhuA2* Δ*(lac-proAB)* *glnV* *galK16* *galE15* *R(zgb-210::Tn10)*Tet$^{S}$ *endA1* *thi-1* Δ*(hsdS-mcrB)5* | New England Biolabs |
| BL21 | *fhuA2* *[lon]* *ompT* *gal* *[dcm]* Δ*hsdS* | New England Biolabs |

**Supplementary Table 2. Plasmids used in this study**

| Name | Description | Reference |
|---|---|---|
| pBJ114 | *galK* containing vector for generation of in-frame deletions in *M. xanthus*, $Kan^R$ | [3] |
| pSW105 | $P_{pilA}$ , $Kan^R$, *attP* | [4] |
| pMal-p5x | Expression vector for periplasmic MalE fusions | NEB |
| pET24b+ | Expression vector for $His_6$-tagged protein | Novagen |
| pMAT150 | pBJ114, in-frame deletion of *pilT* | [1] |
| pBJd*aglQ* | pBJ114, in-frame deletion of *aglQ* | [5] |
| pMH12 | pBJ114, endogenous *tfcP*-sfGFP | This study |
| pMAT167 | pBJ114, in-frame deletion of *pilX1* | [1] |
| pMAT164 | pBJ114, in-frame deletion of *pilY1.1* | [1] |
| pMAT163 | pBJ114, in-frame deletion of *pilB* | [1] |
| pMAT162 | pBJ114, in-frame deletion of *pilA* | [1] |
| pMAT170 | pBJ114, in-frame deletion of *pilB* and *pilT* | This study |
| pMAT336 | pBJ114, endogenous mCherry-PilM | [1] |
| pMAT407 | pSW105, $P_{pilA}$ -*tfcP*-FLAG | This study |
| pMAT409 | pSW105, $P_{pilA}$ -$tfcP^{C91H}$-FLAG | This study |
| pMAT408 | pSW105, $P_{pilA}$ -$tfcP^{C91M}$-FLAG | This study |
| pMAT410 | pSW105, $P_{pilA}$ -$tfcP^{\Delta 118\text{-}153}$-FLAG | This study |
| pMAT210 | pSW105, $P_{pilA}$ -*pilY1.1* | This study |
| pMAT220 | pSW105, $P_{pilA}$ -*fimU1* | This study |
| pMAT222 | pSW105, $P_{pilA}$ -*pilV1* | This study |
| pMAT310 | pSW105, $P_{pilA}$ -*pilX1* | This study |
| pMH45 | pSW105, $P_{pilA}$ -$tfcP^{C31A,C34A}$-FLAG | This study |
| pMAT221 | pSW105, $P_{pilA}$ -*pilW1* | This study |
| pMH60 | pBJ114, endogenous $pilY1.1^{D1173A}$ | This study |
| pMH1 | pBJ114, in-frame deletion of *tfcP* | This study |
| pMH2 | pBJ114, in-frame deletion of *fimU1* | This study |
| pMH3 | pBJ114, in-frame deletion of *pilW1* | This study |
| pMH4 | pBJ114, in-frame deletion of *pilV1* | This study |
| pMH5 | pET24b+, $tfcP\text{-}His_6$ | This study |
| pMH7 | pSW105, $P_{pilA}$ -*tfcP* | This study |
| pMH10 | pBJ114, endogenous *tfcP*-FLAG | This study |
| pMH31 | pMAL-p5x, MalE-TfcP | This study |
| pMH39 | pMAL-p5x, MalE-$TfcP^{C91M}$ | This study |
| pMH45 | pSW105, $tfcP^{C31A,C34A}$-FLAG | This study |
| pEC86 | Constitutive expression of *ccm* genes of *E. coli* | [6] |

### Supplementary Table 3. Oligonucleotides used in this study

| | Oligonucleotides used for cloning | |
|---|---|---|
| **Name** | **Sequence[1]** | **Brief description** |
| 0359-A-HindIII | **GCGCAAGCTT**GCATGGTGACGCTGAGTCCC | Δ*fimU1* |
| 0359-B-XbaI | **GCGCTCTAGAT**CCGCGTGTGTGCCTCATG | Δ*fimU1* |
| 0359-C-XbaI | **GCGCTCTAGAT**GAGCACTGCCGGCACCTGAAG | Δ*fimU1* |
| 0359-D-BamHI | **GCGCGGATCC**CGGAGGTGGAGCTGCTGC | Δ*fimU1* |
| 0360-A-HindIII | **GCGCAAGCTT**AAGGTCTACGCGACCACGGC | Δ*pilW1* |
| 0360-B-XbaI | **GCGCTCTAGA**CGTCTTCACGGCGCCATCCT | Δ*pilW1* |
| 0360-C-XbaI | **GCGCTCTAGA**ACGGAAAATTGAGCATGAGG | Δ*pilW1* |
| 0360-D-BamHI | **GCGCGGATCC**GGAAGTGGCGCAGGCCTTCG | Δ*pilW1* |
| 0361-A-HindIII | **GCGCAAGCTT**CACGGGCTCTGGCATCGCCG | Δ*pilV1* |
| 0361-B-XbaI | **GCGCTCTAGA**CGCTGTCACTGCGGCATCCT | Δ*pilV1* |
| 0361-C-XbaI | **GCGCTCTAGA**ATGGCGCCGTGAAGACGACT | Δ*pilV1* |
| 0361-D-BamHI | **GCGCGGATCC**CAGGTACTCCAGCGTCGGTA | Δ*pilV1* |
| 0363-A-HindIII | **GCGCAAGCTT**GTGCCGCCGCTCAGGCATG | Δ*tfcP* |
| 0363-Bflag-KpnI | **GCGCGGTACC**CTTCTTCCCCTGCGAACG | *tfcP-FLAG* |
| 0363-Cflag-KpnI | **GCGCGGTACC**CTTCTTCCCCTGCGAACG | *tfcP-FLAG* |
| 0363-B-XbaI | **GCGCTCTAGA**GATGAGTCGGTTCATGGG | Δ*tfcP* |
| 0363-C-XbaI | **GCGCTCTAGA**TCGCAGGGGAAGAAGTGA | Δ*tfcP* |
| 0363-D-BamHI | **GCGCGGATCC**GCGACAGGTTTCCGTAGG | Δ*tfcP* |
| sfGFP-B overlay | GCGCGGATGAGGGTGCGCATCATTTGTAGAGCTC | *tfcP-sfGFP* |
| 0363-C overlay | TGATGCGCACCCTCATCCAGACACTGGCCG | *tfcP-sfGFP* |
| 0363 AvaI FactorXa - SP | **GCGCCTCGGG**ATCGAGGGAAGGACGGATGAAGGCAAGCTCGCCTTC | MalE-TfcP/MalE-TfcP$^{C91M}$ |
| 0363 Stop HindIII | **CGCGAAGCTT**TCACTTCTTCCCCTGCGAACG | MalE-TfcP/MalE-TfcP$^{C91M}$ |
| MalE start NdeI* | **GCGCCATATG**AAAATAAAAACAGGTGCACGC | MalE-TfcP/MalE-TfcP$^{C91M}$ |
| 0363 Start XbaI | **GCGCTCTAGA**AACCGACTCATCCTGTTG | PpilA-tfcP |

| | | |
|---|---|---|
| 0363 nostop BamHI | **GCGC<u>GGATCC</u>**CTTCTTCCCCTGCGAACG | $P_{pilA}$-*tfcP*-FLAG/sfGFP |
| 0363 Δ118-153 BamHI | **GCGC<u>GGATCC</u>**AGGAGGTGTGGGGTGGAGGCT | $P_{pilA}$-*tfcP*$^{\Delta 118\text{-}153}$-FLAG |
| 0363-Bmut-XmaI | **GCGC<u>CCCGGG</u>**CGGCGGCCTTCTCGAAGGCGAG | *tfcP*$^{C31A,C34A}$ |
| 0363-Cmut-XmaI | **GCGC<u>CCCGGG</u>**CTCACGTCGTCACCGCGCAAG | *tfcP*$^{C31A,C34A}$ |
| 0363-Short-B | **GCGC<u>TCTAGAG</u>**TGGAGGCTGAGCGCCAG | *tfcP*$^{\Delta 118\text{-}153}$ |
| 0363-Short-C | **GCGC<u>TCTAGA</u>**TGATGCGCACCCTCATCCAGAC | *tfcP*$^{\Delta 118\text{-}153}$ |
| PilY1_1 mut fwd HindIII | **GCGC<u>AAGCTT</u>**CAATCAAAACCAGATCAACAG | *pilY1.1*$^{D1173A}$ |
| PilY1_1_mut rev BamHI | **GCGC<u>GGATCC</u>**GATGAACAAGTGATTGTCATG | *pilY1.1*$^{D1173A}$ |
| 0363-E | GTCTCTTGAGACCAACC | Δ*tfcP* |
| 0363-F | GTCGTAGGGGGAGATTC | Δ*tfcP* |
| 0363-G | CACGGATGAAGGCAAGC | Δ*tfcP* |
| 0363-H | GACGGTTCATCCGCCTG | Δ*tfcP* |
| 0361-0359-E | GCTCACCGGCTGGCGCCATG | Δ*fimU1/pilV1/pilW1* |
| 0361-0359-F | CTTCGACCCGGCGAAGCACG | Δ*fimU1/pilV1/pilW1* |
| 0361-0359-G | CATCGTCTTCAGTGACACGC | Δ*pilW1* |
| 0361-0359-H | GGCGCGACAAGTTCATTGGG | Δ*pilW1* |
| 0363 C91MC | **GCGC<u>ACCGGT</u>**ATGGATACGCGC | *tfcP*$^{C91M}$ |
| 0363 C91HC | **GCGC<u>ACCGGT</u>**CATGATACGCGCCTGC | *tfcP*$^{C91H}$ |
| 0363 C91XB | **GCGC<u>ACCGGT</u>**CTTGGGTTTGATCTGG | *tfcP*$^{C91M/H}$ |
| pilY1.1$^{D1173A+}$ | CGAGACGGCAACTACGCCGTCATGTACGTGCCG | *pilY1.1*$^{D1173A}$ |
| pilY1.1$^{D1173A-}$ | CGGCACGTACATGACGGCGTAGTTGCCGTCTCG | *pilY1.1*$^{D1173A}$ |
| 0361-$G_n$ | CCACCATGGCCATCCTGCTG | Δ*pilV1* |
| 0361-$H_n$ | CAGCTCAGGACGACGCGTAC | Δ*pilV1* |
| 0359-G | CGGTGGCCATCGCCTCCATC | Δ*fimU1* |
| 0359-$H_n$ | GATGGCCTGGTTCTGGGTCG | Δ*fimU1* |
| 0359 start XbaI | **GCGC<u>TCTAGA</u>**ATGAGGCACACACGCGGAATC | $P_{pilA}$-*fimU1* |
| 0359 stop HindIII | **GCGC<u>AAGCTT</u>**TCAGTTCACGCACTCGATGGC | $P_{pilA}$-*fimU1* |
| 0360 start XbaI | **GCGC<u>TCTAGA</u>**GTGAAGACGACTTTGACGC | $P_{pilA}$–*pilW1* |

| | | |
|---|---|---|
| 0360 stop HindIII | **GCGCAAGCTT**TCAATTTTCCGTCAGGAG | $P_{pilA}$–*pilW1* |
| 0361 start XbaI | **GCGCTCTAGA**GTGAAGACGACTTTGACGCG | $P_{pilA}$-*pilV1* |
| 0361 stop HindIII | **GCGCAAGCTT**TCACGGCGCCATCCTCGTC | $P_{pilA}$-*pilV1* |
| 0362 start XbaI | **GCGCTCTAGA**GTGATGCGCACCCTCATCCAG | $P_{pilA}$-*pilY1.1* |
| 0362 stop HindIII | **GCGCAAGCTT**TCACTGCGGCATCCTCCCGTC | $P_{pilA}$-*pilY1.1* |
| 0364 start XbaI | **GCGCTCTAGA**GTGCAACGTCCCACAACC | $P_{pilA}$–*pilX1* |
| 0364 stop HindIII | **GCGCAAGCTT**TCAAGGGGCGGGGGTGATG | $P_{pilA}$–*pilX1* |
| | **Oligonucleotides used for operon mapping** | |
| **Name** | **Sequence** | **Combination** |
| 0365 map-fwd | CCGAGCCATCCGAGGTG | A |
| pilX1 map-rev | GTGGGACGTTGCACCATGT | A |
| pilX1 q-1 fwd | AGGTCTCGACGATGACAATGG | B |
| tfcP q-1 rev | CCGCGGCCTTTGTTTTCTTC | B |
| tfcP q-2 fwd | ACAGAAGAAAACAAAGGCCGC | C |
| PilY1 map rev | CGGTGATTCCTCGGTGATG | C |
| PilY1 map fwd | CTGAGCCAGGACGAGAGCG | D |
| PilV map rev | AGCGTGATGTTCTCCATCGC | D |
| pilV1 q-2 fwd | CCCTCCATCCTCAGCACTATC | E |
| PilW map rev | TGAAGACGATGGGGGCATTC | E |
| pilW1 q-1 fwd | ATGTCGAACGTCTCTCGGTG | F |
| fimU1 q-1 rev | CATTCTCGCGTTGACGGTTG | F |
| | **Oligonucleotides used for qRT-PCR** | |
| **Name** | **Sequence** | **Gene** |
| fimU1_q-2_fwd | CTCCGGTGCGACAATGAATG | *fimU1* |
| fimU1_q-2_rev | TGTAGCAGAGCCCGTGAATC | *fimU1* |
| pilV1_q-1_fwd | GGACTGGATGAGAGCTACGTC | *pilV1* |
| pilV1_q-1_rev | GCTCGATAGTGCTGAGGATGG | *pilV1* |
| pilW1_q-1_fwd | ATGTCGAACGTCTCTCGGTG | *pilW1* |
| pilW1_q-1_rev | CGACAAGTTCATTGGGGTGC | *pilW1* |

| | | |
|---|---|---|
| pilY1.1_q-2_fwd | GACGTCTCCCATTACGACCC | *pilY1.1* |
| pilY1.1_q-2_rev | AAGATGACTTCCTTCCCGCC | *pilY1.1* |
| rpsS_q-1_rev | GACGAACACCGGGATGAACT | *rpsS* |
| rpsS_q-2_fwd | GTTCGATCAAGAAGGGTCCGT | *rpsS* |
| tfcP_q-1_fwd | CCGACTCATCCTGTTGTCCC | *tfcP* |
| tfcP_q-1_rev | CCGCGGCCTTTGTTTTCTTC | *tfcP* |
| pilA_q-1_fwd | GATTCAACCCCCGCAACCG | *pilA* |
| pilA_q-1_rev | GTTCGTCTTCGCCTCGGAC | *pilA* |
| PilX1_q-6_fwd | GGTCGGAGGCTGGAACTC | *pilX1* |
| PilX1_q-6_rev | TCGTTTGGAGCGGGAAGG | *pilX1* |
| Tau_q-3_fwd | AGTGGAAGTCGTTGGTCTGC | *tuf2 (mxan_3298)* |
| Tau_q-3_rev | TTGGTGTGCGGGGTGATG | *tuf2 (mxan_3298)* |

[1]Restriction sites are underlined. Oligonucleotide sequences that are not complementary to the template are indicated in bold.

**Supplementary Table 4. Heavy labelled ($^{13}$C and $^{15}$N) reference peptides with C-terminal Lys or Arg residue**

| Peptide Name | Peptide Sequence |
|---|---|
| FimU1_1 | DFLDDLPALDAAAPGNLR |
| FimU1_2 | IVVEENVPR |
| FimU1_3 | SLIQVEPR |
| PilA_1 | FGANSAIDDPTPVVAR |
| PilA_2 | NAADLPVPAAGVPCISNDSFR |
| PilA_3 | VSAAAGDCEVR |
| PilA_4 | YSDFANEIGFAPER |
| PilB_1 | ENLISVQQLR |
| PilB_2 | HLVVPVNR |
| PilB_3 | LGMSSLR |
| PilC_1 | DILVFTR |
| PilC_2 | KGEMEAMDVEAVNAR |
| PilC_3 | TLGTMISSGVPILDALDVTAK |
| PilC_4 | TVEDAIIYVR |
| PilM_1 | DVTIGGNQFTEEIQK |
| PilM_2 | QLNVSYEEAEALK |
| PilM_3 | SLDFYAGTAADSNFSK |
| PilM_4 | VLSSVAEQVAGEIQR |
| PilN_1 | INLLPVR |
| PilN_2 | LAVLDALR |
| PilN_3 | MMDALASATPK |
| PilN_4 | QSELEAHQAGVASTK |
| PilN_5 | QVGGAQVGVPILVEFK |
| PilO_1 | DIEELLAQINDIGKK |
| PilO_2 | LSEALTELPEQR |
| PilO_3 | VVLQSEFQATTFR |
| PilP_1 | LVAVVTGDASPVAMVEDPAGR |
| PilP_2 | QDPAYNMMTGR |
| PilQ_1 | ALGKEEFGNIIR |
| PilQ_2 | NIVVADDVSGK |
| PilQ_3 | TNVLIVK |
| PilT_1 | GASDLHVTTGSPPQLR |
| PilT_2 | VHQIYSSMQVGQAK |
| PilV1_1 | DLVPGVPDTAGNIANVR |
| PilW1_1 | AGSGMGNAPIVFSDTR |
| PilW1_2 | ALFEEQTMLAQVTGR |
| PilW1_3 | INVVPGTGIETTTTDR |
| PilW1_4 | LQPTTAPTTPALLVNPAR |
| PilW1_5 | NLACHVEVTNVDAAGR |
| PilX1_1 | QSPSGDAYAAFPLQTNVR |

| | |
|---|---|
| PilX1_2 | YKEAYFAAEAGLAEGR |
| PilY1.1_1 | SATVSGDLSPDIANDFVITK |
| PilY1.1_3 | SSNIEHAFSTAK |
| PilY1.1_4 | VNLDQVNPNAPLGQK |
| TfcP_1 | AWLAGPNQIKPK |
| TfcP_2 | GPSVDLGPVVPMR |
| TsaP_1 | GDLVGPVGER |
| TsaP_2 | IGVDLANSVPVTTQGFVTQR |
| TsaP_3 | SLEELVPGDR |
| TsaP_4 | YVVYHTTQAVK |

### Supplementary References


1 Treuner-Lange, A. *et al.* PilY1 and minor pilins form a complex priming the type IVa pilus in *Myxococcus xanthus*. *Nat Commun* **11**, 5054 (2020).

2 Friedrich, C., Bulyha, I. & Søgaard-Andersen, L. Outside-in assembly pathway of the type IV pilus system in *Myxococcus xanthus*. *J Bacteriol* **196**, 378-390 (2014).

3 Julien, B., Kaiser, A. D. & Garza, A. Spatial control of cell differentiation in *Myxococcus xanthus*. *Proc Natl Acad Sci U S A* **97**, 9098-9103 (2000).

4 Jakovljevic, V., Leonardy, S., Hoppert, M. & Søgaard-Andersen, L. PilB and PilT are ATPases acting antagonistically in type IV pilus function in *Myxococcus xanthus*. *J Bacteriol* **190**, 2411-2421 (2008).

5 Sun, M., Wartel, M., Cascales, E., Shaevitz, J. W. & Mignot, T. Motor-driven intracellular transport powers bacterial gliding motility. *Proc Natl Acad Sci U S A* **108**, 7559-7564 (2011).

6 Arslan, E., Schulz, H., Zufferey, R., Kunzler, P. & Thöny-Meyer, L. Overproduction of the *Bradyrhizobium japonicum* c-type cytochrome subunits of the cbb3 oxidase in *Escherichia coli*. *Biochem Biophys Res Commun* **251**, 744-747 (1998).
